# CDK4 inactivation balances resistance to apoptosis with heightened metabolic sensitivity in triple negative breast cancer cells

**DOI:** 10.1101/2024.09.17.613455

**Authors:** Dorian V. Ziegler, Lucia Leal-Esteban, Kanishka Parashar, Wilson Castro, Nadège Zanou, Katharina Huber, Jaime López-Alcalá, Xavier Pascal Berney, María M Malagón, Catherine Roger, Marie-Agnès Berger, Giulia Paone, Hector Gallart-Ayala, Julijana Ivanisevic, Jennifer Rieusset, Melita Irving, Lluis Fajas

## Abstract

The shift in the energetic demands of proliferating cells during tumorigenesis requires intense crosstalk between the cell cycle and metabolism. Beyond their role in cell proliferation, cell cycle regulators also modulate intracellular metabolism in normal tissues. However, in the context of cancer, where CDK4 is upregulated or stabilized, the metabolic role of CDK4 is barely understood. Using both genetic and pharmacological approaches, we aimed to determine the metabolic role of CDK4 in TNBC cells. Unexpectedly, deletion of CDK4 only slightly reduced triple-negative breast cancer (TNBC) cell proliferation and allowed tumor formation *in vivo*. Furthermore, proapoptotic stimuli failed to induce appropriate cell death in TNBC cells with CDK4 depletion or long-term CDK4/6 inhibitor treatment. Mechanistically, CDK4 enhances mitochondria-ER contact (MERC) formation, thus promoting mitochondrial fission and ER-mitochondrial calcium signaling. Phosphoproteomic analysis also revealed a role for CDK4 in regulating PKA activity at MERCs to sustain ER-mitochondrial calcium signaling. This proper CDK4-mediated mitochondrial calcium signaling is then required for metabolic flexibility of TNBC cells. Taken together, these results demonstrate that CDK4 inhibition leads to cell death resistance, inhibiting mitochondrial apoptosis and functions through attenuated MERCs formation and ER-mitochondrial calcium signaling in TNBC. Overall, this study provides new insights into the mechanisms of TNBC resistance to CDK4/6i therapy and paves the way to explore potential synergistic therapeutic targeting MERCs-associated metabolic shifts.

**Graphical Abstract:** 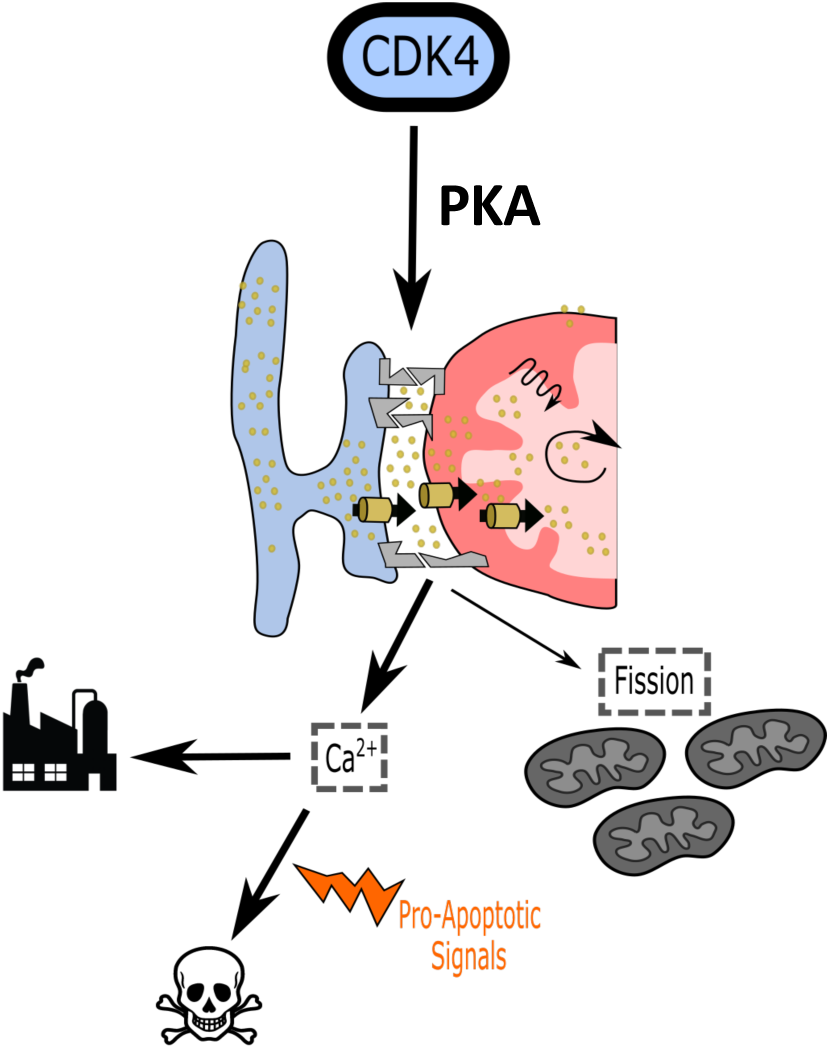

## Introduction

Cell cycle regulators are key factors for cell proliferation and participate in a myriad of pathophysiological processes, including development, tissue regeneration, and cancer^1^. In dividing cells, entry into S-phase depends on the activation of G1 Cyclins/Cyclin-Dependent Kinases (CDKs) 4 and 6, which phosphorylate the retinoblastoma protein (RB), resulting in the release of the E2F transcription factor to progress into the cell cycle ^2^. As a canonical driver of cell cycle progression, the cyclin/CDK-RB-E2F pathway is commonly deregulated during tumor progression; thus, CDK4/6 inhibitors (CDK4/6i) constitute a valuable antitumor therapeutic approach^3,4^. Unfortunately, in triple-negative breast cancer (TNBC), significant improvements in patients were not observed in clinical trials with CDK4/6i^5^. This outcome revealed the importance of therapeutic resistance in this context, which appears to occur *via* multiple mechanisms and usually results from long-term treatment with CDK4/6i^5^.

In addition to its role in cell cycle regulation, CDK4 has also been shown to be involved in other biological processes. For instance, CDK4 regulates cellular metabolism mainly through the activation of anabolic processes and the repression of oxidative catabolism ^6–8^. Moreover, CDK4 modulates cell fate through the control of not only autophagy but also two antitumorigenic processes, cellular senescence and apoptosis^9–12^. Finally but importantly, CDK4 participates in the modulation of cancer cell dynamics through interactions with cytoskeleton-associated proteins or epithelial–mesenchymal transition (EMT)-promoting factors. Based on this pleiotropic role of CDK4, the effect of treatment with CDK4/6i, even in the short term, may affect other cellular biological processes in addition to the cell cycle.

The early discovery of the Warburg effect pushed mitochondria to the background of cancer biology as a second, even nonexistent, source of energy for cancer cells^13^. However, in the mid-1990s, interest in the importance of mitochondria in cancer was renewed with the demonstration of their role in intrinsic cell death. Regulating multiple prosurvival pathways, including anabolic biosynthesis, calcium signaling and redox balance, mitochondrial reprogramming is currently considered a hallmark of tumor cells^14^. As highly dynamic organelles, mitochondria have been observed to be sensitive to imbalances in dynamics in different cancer contexts^15,16^. Mitochondria-endoplasmic reticulum contacts sites (MERCs), also termed mitochondria-associated membranes (MAMs), regulate mitochondrial function and dynamics under physiological conditions^17^, and their role in tumorigenesis is beginning to be delineated^18–20^. Participating in the regulation of calcium signaling, redox balance, phospholipid synthesis, mitochondrial fission and intrinsic cell death, MERCs are indeed highly modulated in cancer cells, with functions in both cancer bioenergetics or escape from cell death^21^. However, the upstream control of these MERCs in cancer-specific contexts is barely understood. Specifically, the role of cell cycle regulators in the regulation of MERCs is completely unknown.

This study aims to decipher the mechanism by which CDK4 regulates the fate of TNBC cells and tumors in mice. In elucidating the cell cycle-dependent effect of CDK4 using proliferating CDK4-KO cells, we hypothesized that CDK4 controls MERCs, thus functioning at the intersection between the cell cycle machinery, metabolism and intrinsic apoptosis through modulation of calcium signaling.

## Results

### CDK4 is dispensable for tumor growth *in vitro* and *in vivo*

CDK4 is involved in progression into the cell cycle through the regulation of RB phosphorylation and the subsequent release of E2F transcription factors. To discriminate between the cell cycle-mediated and cell cycle-independent effects of CDK4, we used a specific CRISPR CDK4 knockout (KO) model in MDA-MB-231 TNBC cells that we previously developed in the laboratory (Fig 1A)^10^. We revealed that stable CDK4 depletion in these cells resulted in a constitutively high level of RB phosphorylation at serine 780 (S780) (Fig 1A-B). This high level of RB S780 phosphorylation was independent of the stabilization of CDK6 (Fig 1A) but was associated with slightly increased levels of CDK4-associated cyclins, namely, Cyclin D1 and Cyclin D3 (Sup. Fig 1A). Consistent with these results, CDK4 KO in TNBC cells only slightly reduced their proliferation (Fig 1C). In line with the sustained proliferation of CDK4-KO cells, RNA-seq analysis showed only a weak decrease in the expression of cell cycle-related genes and E2F target genes in CDK4-depleted cells (Sup. Fig 1B). Interestingly, short-term treatment of CDK4-KO cells with the CDK4/6 inhibitor abemaciclib reduced RB S780 phosphorylation, whereas long-term chemical inhibition did not (Fig. 1E-F). Short-term CDK4/6 inhibition thus reduced the early proliferation of TNBC cells, while long-term inhibition resulted in the generation of resistant and proliferating cells after a few passages (Fig 1G). The effects of long-term CDK4/6 inhibition were, therefore, reminiscent of those of CDK4 KO in TNBC cells.

**Figure 1:**
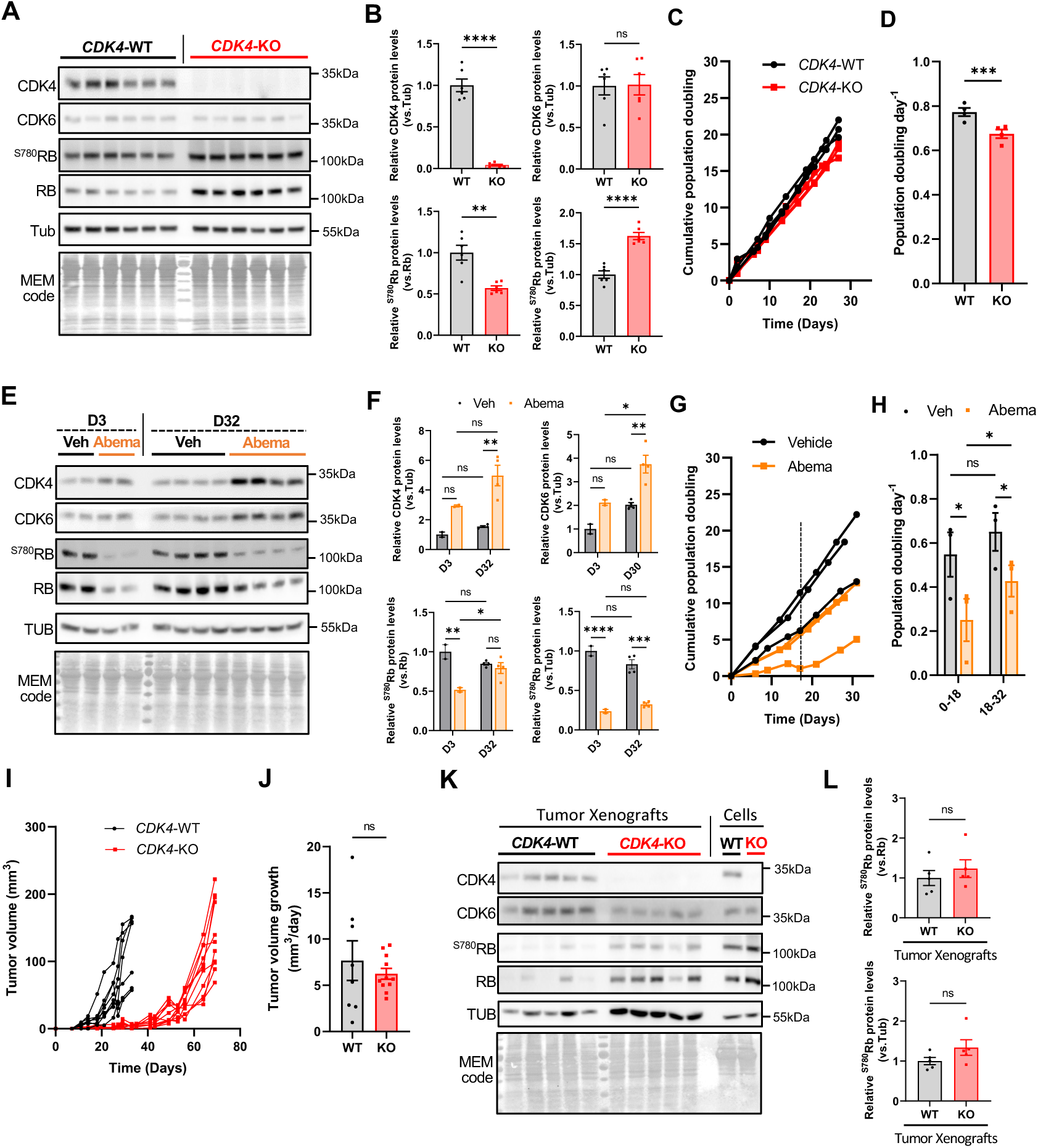
CDK4 is dispensable for TNBC tumor growth *in vitro* and *in vivo*. **(A-B).** Immunoblots and relative protein levels of CDK4, CDK6, ^S780RB^, RB, Tubulin (TUB) and MEM code of CDK4-WT and—KO TNBC cells. N=6 independent biological replicates. Unpaired T-tests. **(C-D.)** Growth curves and proliferation rates of CDK4-WT and—KO TNBC cells. N=3 independent biological replicates. Paired T-test. **(E-F.)** Immunoblots and relative protein levels of CDK4, CDK6, ^S780RB^, RB, Tubulin (TUB) and MEM code of TNBC cells, upon treatment with Vehicle (Veh) or Abemaciclib (Abema) for 3 days (D3) or 32 days (D32). N=2-4 independent biological replicates. 2-way ANOVA and Tukey’s multiple comparison tests. **(G-H.)** Growth curves and proliferation rates of TNBC cells, upon treatment with Vehicle (Veh) or Abemaciclib (Abema) N=3 independent biological replicates. 2-way ANOVA and Tukey’s multiple comparison tests. **(I).** Tumor volume of CDK4-WT and—KO tumor xenografts. One curve represents one xenograft. N=8 WT and N=10 KO. **(J).** Tumor volume growth after tumors reached 20 mm ^3^ of volume of CDK4-WT and—KO xenografts. N=8 WT and N=10 KO. Welch’s T-test. **(K-L).** Immunoblots and relative protein levels of CDK4, CDK6, ^S780RB^, RB, Tubulin (TUB) and MEM code of CDK4-WT and—KO tumor xenografts, and CDK4-WT and—KO TNBC cells. N=5 WT and KO tumors. Unpaired T-tests. ns: nonsignificant/*p-value*> 0,1,* *p-value* <0,05, ** *p-value* <0,01, *** *p-value* <0,001, **** *p-value* <0.0001.

Since CDK4-KO MDA-MD-231 cells were still able to proliferate, we next explored the capacity of these cells to form tumors in mice when orthotopically implanted into the mammary fat pads of immunocompromised mice. Strikingly, these CDK4-KO cells were able to form tumors, with the same penetrance as CDK4-WT cells, but with a latency of 21 days (Fig 1I) (Sup. Fig 1C). After this period, however, a similar tumor growth rate (Fig 1J) and an equivalent tumor volume at sacrifice were observed between the CDK4-WT and CDK4-KO groups (Sup. Fig 1C-D). In line with the *in vitro* results, CDK4-KO tumors did not show changes in RB S780 phosphorylation (Fig 1K-L). Finally, the expression of the proliferation marker Ki67 was unchanged in CDK4-KO tumors compared to CDK4-WT tumors (Sup. Fig 1E).

Collectively, these results indicated that CDK4 is dispensable for cell cycle progression in TNBC cells, which validates the appropriateness of this model for exploring the cell cycle-independent functions of CDK4.

### CDK4 facilitates apoptosis in TNBC cells

By analysis of cohort databases, we observed that in a cohort of 256 chemotherapy-treated TNBC patients, high expression of CDK4 was, surprisingly, associated with a better prognosis than low expression of CDK4 (Sup. Fig 2A). These clinical data suggested that CDK4 could mediate the chemotherapy response of TNBC cells. We therefore investigated whether CDK4-KO cells are affected by proapoptotic stresses, including not only the stress imposed by the chemotherapeutic drug cisplatin but also oxidative stress (H_2_0_2_) and mitochondrial stress (oligomycin and antimycin A) (O+A). Upon exposure to these three proapoptotic stresses, CDK4-KO cells exhibited greater resistance than the control CDK4-WT cells, as evidenced by the increased cell number and cell viability after treatment (Fig 2A-B and Sup. Fig 2B). This increased viability of CDK4-KO cells was also associated with failure to induce cleavage of caspase 3, a marker of apoptosis, upon treatment (Fig 2C). Independent clones with CRISPR‒ Cas9-mediated KO of CDK4 were also resistant to apoptosis induced by H_2_0_2_ or O+A (Sup. Fig 2C-D). Reintroduction of CDK4 into the KO cells restored the apoptotic phenotype in response to the same proapoptotic stimuli (Sup. Fig 1E-G), indicating that the observed effects were caused by CDK4. We also investigated whether CKD4/6i may impact apoptosis sensitivity. Consistent with the results in CDK4-KO cells, long-term abemaciclib pretreatment was also sufficient to induce resistance to cisplatin and other proapoptotic stimuli, as evidenced by the increased number of viable cells but the reduction in Caspase-3 cleavage upon treatment (Fig 2D-F).

**Figure 2:**
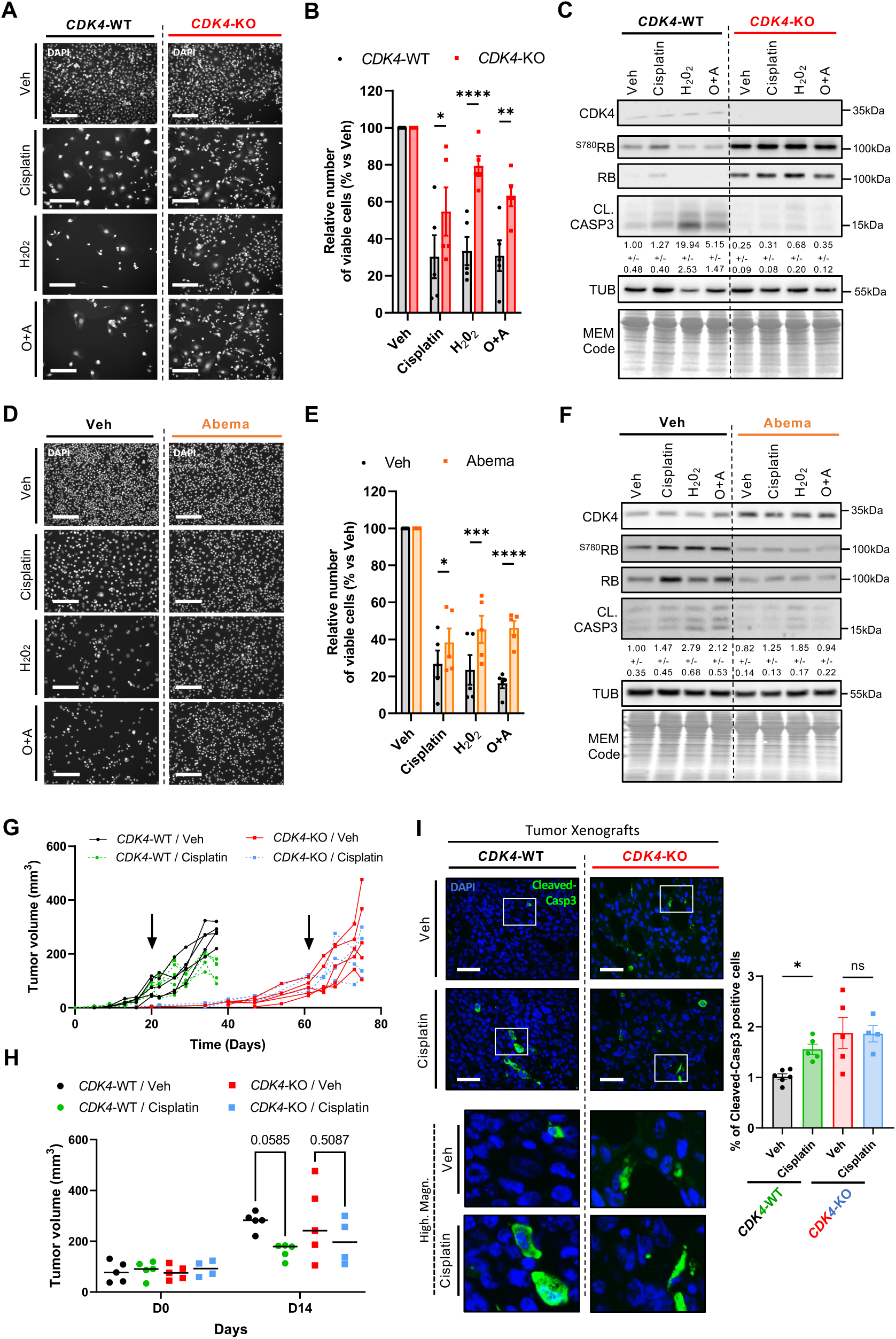
CDK4 facilitates apoptosis in TNBC. **(A-B).** Representative pictures of DAPI staining and quantification of the number of viable CDK4-WT and—KO TNBC cells, upon treatment with Vehicle, Cisplatin, H_2_0_2_, or Oligomycin and Antimycin (O+A). Scale bars: 200 μm. N=5 independent biological replicates. 2-way ANOVA and Sidák’s multiple comparison tests. **(C).** Immunoblots and relative protein levels of CDK4, ^S780RB^, RB, Cleaved Caspase-3 (CL.CASP3), Tubulin (TUB) and MEM code of CDK4-WT and—KO TNBC cells, upon treatment with Vehicle, Cisplatin, H_2_0_2_, or Oligomycin and Antimycin (O+A). Quantification of relative cleaved Caspase-3 protein levels (normalized to tubulin level). Mean +/—N=3 independent biological replicates. **(D-E.)** Representative pictures of DAPI staining and quantification of the number of viable pretreated cells with Vehicle or Abemaciclib (Abema) for 8 days and after treatment with Vehicle, Cisplatin, H_2_0_2_, or Oligomycin and Antimycin (O+A). Scale bars: 400 μm. N=5 independent biological replicates. 2-way ANOVA and Sidák’s multiple comparison tests. **(F).** Immunoblots and relative protein levels of CDK4, ^S780RB^, RB, Cleaved Caspase-3 (CL.CASP3), Tubulin (TUB) and MEM code of pretreated cells with Vehicle or Abemaciclib (Abema) for 8 days and after treatment with Vehicle, Cisplatin, H_2_0_2_, or Oligomycin and Antimycin (O+A). Quantification of relative cleaved Caspase-3 protein levels (normalized to tubulin level). Mean +/—N=3 independent biological replicates. **(G).** Tumor volume of CDK4-WT and—KO tumor xenografts. Arrows indicate when tumors reached 80mm^3^ and the starting point of vehicle (Veh) and Cisplatin treatments. One curve represents one tumor xenograft. n=5 WT Veh, N=5 WT Cisplatin, N=5 KO Veh and N=4 KO Cisplatin. **(H).** Tumor volume of CDK4-WT and—KO tumor xenografts at the end of the sacrifice. 2-way ANOVA and Sidák’s multiple comparison tests**. (I).** Immunofluorescence of slices of CDK4-WT and—KO xenografts for Cleaved-Caspase 3 (green) and DAPI (blue). Representative pictures and associated quantification of Cleaved-Caspase 3 positive cells. 2-way ANOVA and Tukey’s multiple comparison tests. ns: nonsignificant/*p-value*> 0,1, * *p-value* <0,05, ** *p-value* <0,01, *** *p-value* <0.001.

Using an orthotopic xenograft model of human breast cancer, we then evaluated the tumorigenic potential of CDK4-KO cells upon cisplatin treatment *in vivo*. Similar to CDK4-WT cells, CDK4-KO cells formed tumors in engrafted mice, although with a latency period of three weeks. When the average volume of either the CDK4-WT or CDK4-KO tumors reached 80 mm^3^, the mice were treated with cisplatin (Sup. Fig 2H). Cisplatin treatment effectively decreased the growth of CDK4-WT xenograft tumors (Fig 2G-H and Sup. Fig 2I-J). However, the same cisplatin treatment regimen did not reduce tumor growth as effectively in CDK4-KO xenograft tumors (Fig 2G-H and Sup. Fig 2I-J). Accordingly, the induction of caspase 3 cleavage observed in CDK4-WT xenograft tumors upon cisplatin treatment was blunted in CDK4-KO tumors (Fig 2I).

### CDK4 participates in mitochondrial fission

An important function of mitochondria, in addition to the control of bioenergetics, is the regulation of apoptosis in response to specific challenges. To address whether mitochondrial changes could underlie the resistance to apoptosis in CDK4-KO cells (Fig 2), we characterized mitochondrial biology in these cells. Using transmission electron microscopy (TEM), we revealed that CDK4-KO cells exhibit enlarged mitochondria, as evidenced by the increased individual mitochondrial area (Fig 3A-B) and the appearance of elongated giant mitochondria (Fig 3C and Sup. Fig 3A). To a lesser extent, increased mitochondria area was also observed in CDK4-KO xenograft tumors (Sup. Fig 3B). Interestingly, the total mitochondrial number was not affected by the loss of CDK4 (Sup. Fig 3C). Furthermore, Peroxisome Proliferator-Activated Receptor Gamma Coactivator 1-Alpha (PGC1-α) and Estrogen Related Receptor Alpha (ERRα)-target genes were not enriched in CDK4-KO cells, as determined by gene set enrichment analysis (GSEA) of the RNA sequencing (RNA-seq) data (Sup. Fig 3D), suggesting that these giant mitochondria were not the consequence of increased mitochondrial biogenesis. Consistent with this hypothesis, the mRNA levels of genes involved in mitochondrial biogenesis, namely, *Polmrt, Tbf1m* and *Tbf2m* (Sup. Fig 3E) were similar between CDK4-WT and CDK4-KO TNBC cells, indicating that the mitochondrial biogenesis transcriptional program is not activated in CDK4-KO cells.

**Figure 3:**
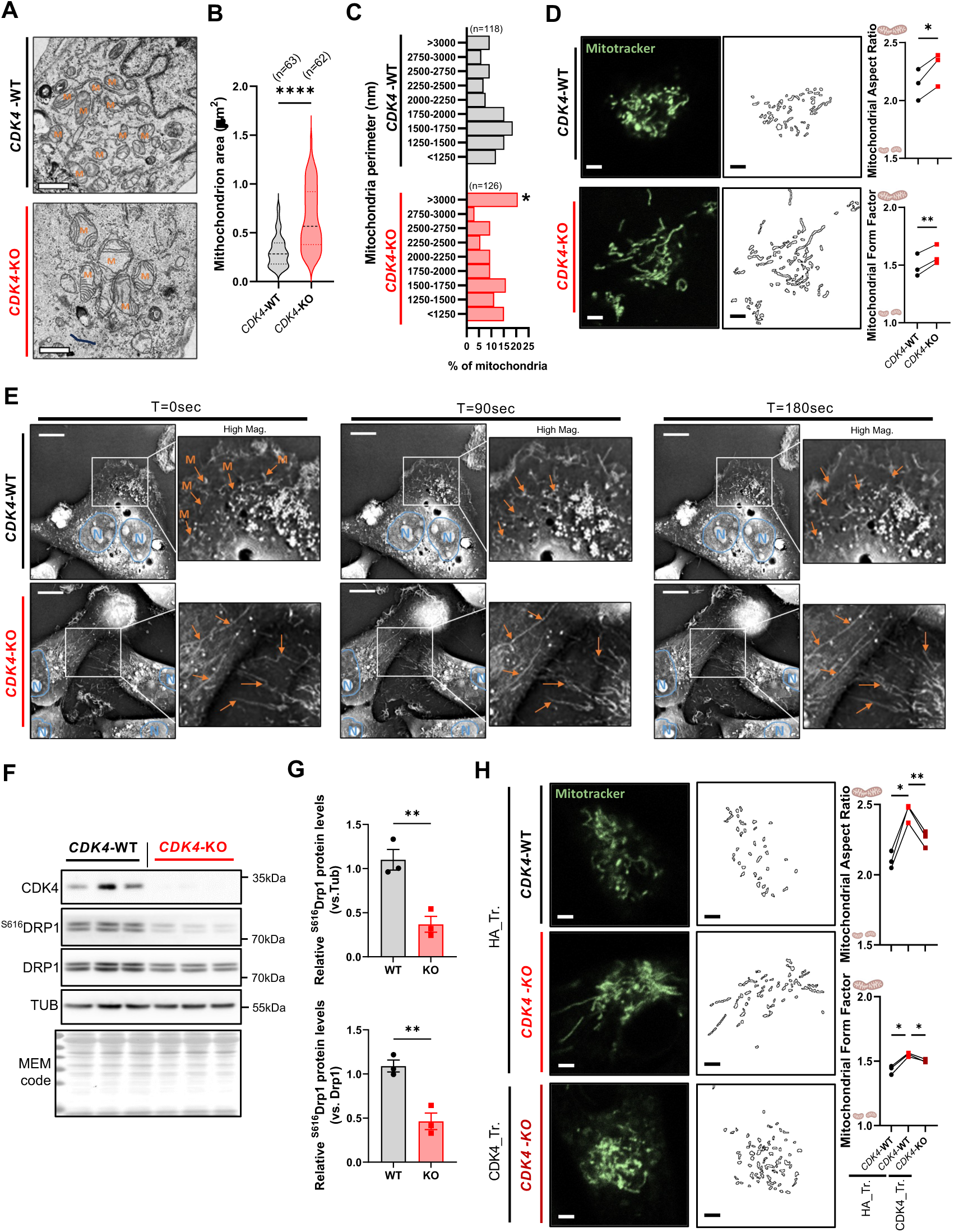
CDK4 participates in mitochondrial fission of TNBC. **(A).** Representative electron micrographs of CDK4-WT and—KO TNBC cells. Scale bars: 1 μM. M=Mitochondria. **(B).** Quantification of mitochondria area of CDK4-WT and—KO TNBC cells. n=mitochondria number representative from N=3 independent biological replicates. Mann-Whitney Test. **(C).** Distribution histogram of mitochondria perimeter of CDK4-WT and—KO TNBC cells. n=mitochondria number representative from N=3 independent biological replicates. Mann-Whitney Test. **(D).** Representative pictures of mitochondria staining using Mitotracker of CDK4—WT and—KO TNBC cells. Scale bars: 4 μM. Associated quantification of mitochondrial aspect ratio (major axis/minor axis) and form factor (1/circularity) in CDK4-WT and—KO TNBC cells. N=3 independent biological replicates, representative of n=85 (CDK4-WT) and n=90 (CDK4-KO) cells. Paired T-tests. **(E).** Representative time-live tracking of mitochondria of CDK4-WT and—KO TNBC cells at T=0sec, T=90sec and T=180sec. N delineation indicates nuclei (Blue) and M arrows indicates mitochondria (Orange). **(F-G).** Immunoblots and relative protein levels of CDK4, ^S616^DRP1, DRP1, Tubulin (TUB) and MEM code of CDK4-WT and—KO TNBC cells. N=3 independent biological replicates. Unpaired T-tests. **(H).** Representative pictures of mitochondria staining using Mitotracker of CDK4-WT, CDK4-KO TNBC cells transfected with empty plasmid HA (HA_Tr.) and CDK4-KO TNBC cells expressing endogenous CDK4 (CDK4_Tr.). Scale bars: 4 uM. Associated quantification of mitochondrial aspect ratio and form factor. N=3 independent biological replicates representing n=148 (CDK4-WT/HA_Tr.), n=151 (CDK4-KO/HA_Tr.) and n=188 (CDK4-KO/CDK4_Tr) cells. RM 1 way ANOVA and Tukey’s multiple comparisons test.

As mitochondria are highly dynamic organelles, we more precisely investigated the mitochondrial shape through evaluation of key mitochondrial morphological parameters, namely, the aspect ratio and form factor. Using fluorescence tracking and electron microscopy, we observed that mitochondria in CDK4-KO cells displayed a higher aspect ratio and form factor (Fig. 3B-C and Sup. Fig 3F), suggesting an increase in elongated mitochondria. Furthermore, dynamic cell imaging showed a persistent fragmented mitochondrial network in CDK4-WT cells and a permanent hyperfused mitochondrial network in CDK4-KO cells (Fig 3E and Sup. Video 1). Mitochondrial dynamics are governed by the fission/fusion equilibrium, which involves coordination between fission proteins (MFF, MIEF and DRP1) and fusion proteins (MFN1, OMA1 and OPA1)^22^ (Giacomello et al. 2020). Since we did not observe significant enrichment of mitochondrial fusion genes based on mRNA sequencing (Sup. Fig 3H), we hypothesized that the appearance of giant mitochondria could be secondary to a defect in the fission process. Indeed, the level of S616-phosphorylated DRP1, a profission protein, was decreased in CDK4-KO cells compared to CDK4-WT cells (Fig 3F-G). Decreased phosphorylation of DRP1 at S616 was also found in TNBC cells treated with CDK4/6i (Sup. Fig 3I-J), accompanied by an increased fused mitochondrial network (Sup. Fig 3K). Finally, we confirmed the role of CDK4 in the mitochondrial fission process by ectopically expressing CDK4, which normalized the mitochondrial aspect ratio and form factor, finally resulting in the restoration of a normal mitochondrial shape in CDK4-KO cells (Fig 3H).

### CDK4 modulates ER-mitochondrial (ER-MT) calcium signaling

Mitochondrial calcium uptake is a determinant of the early steps of apoptosis induction^23^. Indeed, a deficiency in mitochondrial calcium uptake upon proapoptotic stress may allow cells to evade apoptosis^24,25^. Interestingly, we found that CDK4-KO cells exhibited decreased mitochondrial calcium uptake upon exposure to proapoptotic oxidative and mitophagic stresses (Fig 4A-D and Sup. Fig 4A-B).

**Figure 4:**
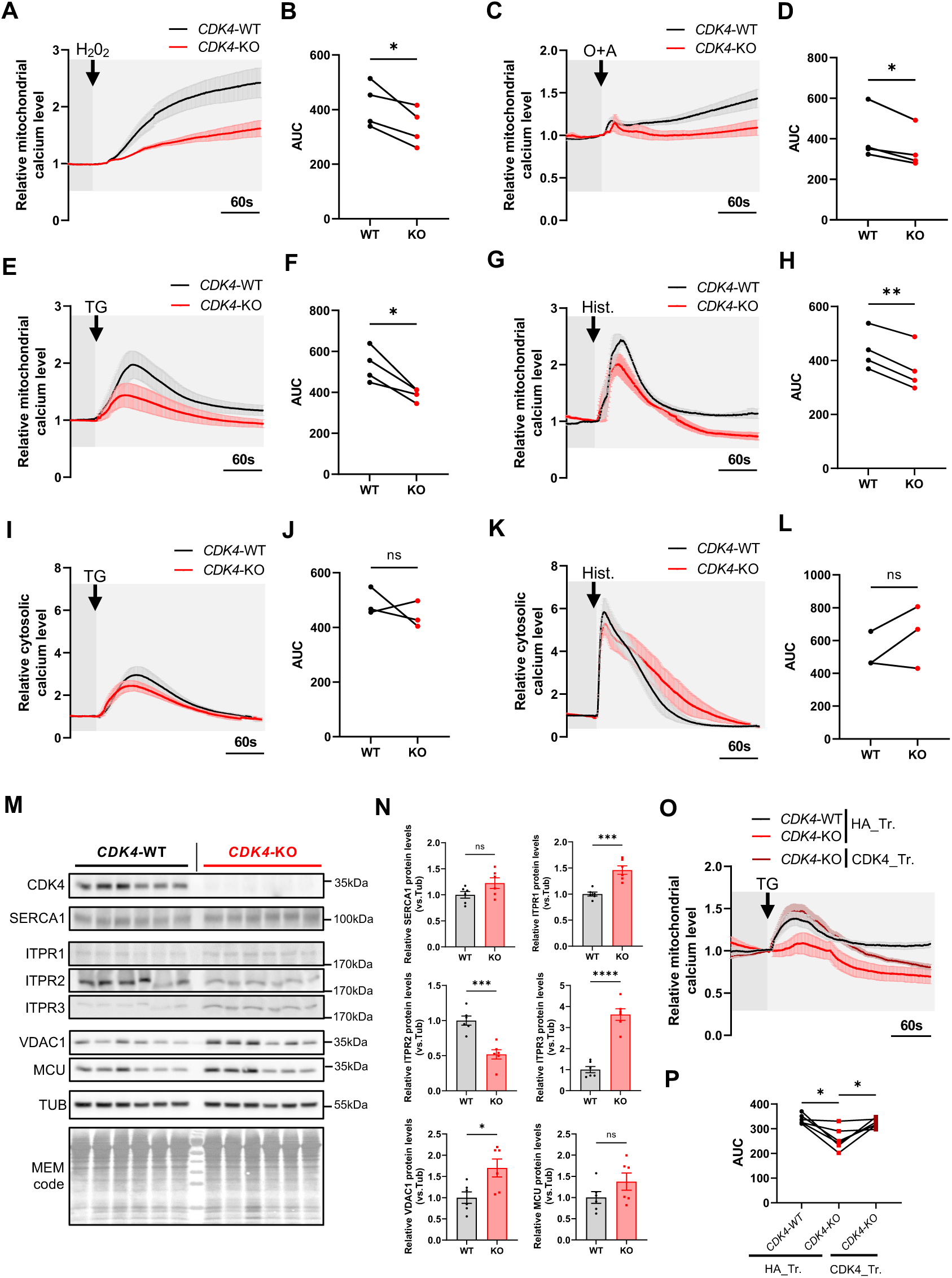
CDK4 regulates ER-mitochondrial calcium signaling. **(A-D).** Relative mitochondrial calcium levels of CDK4-WT and CDK4 KO-TNBC cells upon H_2_0_2_ (2.5 mM) or Oligomycin+Antimycin A (O+A) (100 μM, 10 μM) injections. Curves based on Rhod-2AM probe fluorescence intensity. Arrows indicate the time of injection. Associated quantification of the area under the curves (AUC). N=4 independent biological replicates representing a total of n=135 cells for 10 independent injections (WT-H_2_0_2_), n=123 cells for 9 independent injections (KO-H_2_0_2_), n=128 cells for 9 independent injections (WT-O+A) and n=128 cells for 9 independent injections (KO-O+A). Paired T-tests**. (E-G.)** Relative mitochondrial calcium levels of CDK4-WT and CDK4 KO-TNBC cells upon Thapsigargin (TG) (2 μM) or Histamine (Hist) (50 μM) injections. Curves based on Rhod-2AM fluorescence intensity. Arrows indicate the time of injection. Associated quantification of the area under the curves. N=4 independent biological replicates representing a total of representing a total of n=148 cells for 13 independent injections (WT-TG), n=142 cells for 13 independent injections (KO-TG), n=133 cells for 10 independent injections (WT-Hist.) and n=141 cells for 12 independent injections (KO-Hist.). Paired T-tests. **(I-L.)** Relative cytosolic calcium levels of CDK4-WT and CDK4-KO TNBC cells upon Thapsigargin (TG) (2 μM) or Histamine (Hist) (50 μM) injections. Curves based on Fluo-4 probe fluorescence intensity. Arrows indicate the time of injection. Associated quantification of the area under the curves. N=3 independent biological replicates representing a total of n=107 cells for 9 independent injections (WT-TG), n=123 cells for 9 independent injections (KO-TG), n=121 cells for 9 independent injections (WT-Hist.) and n=118 cells for 9 independent injections (KO-Hist.). Paired T-tests. **(M-N.)** Immunoblots and relative protein levels of CDK4, SERCA1, ITPR1, ITPR2, ITPR3, VDAC1, MCU, Tubulin (TUB) and MEM code of CDK4-WT and—KO TNBC cells. N=6 independent biological replicates. Unpaired T-tests. **(O-P.)** Relative mitochondrial calcium levels of CDK4-WT, CDK4-KO TNBC cells transfected with empty plasmid HA (HA_Tr.) and CDK4-KO TNBC cells expressing endogenous CDK4 (CDK4_Tr.) upon Thapsigargin (TG) (2 μM) injection. Arrows indicate the time of injection. Associated quantification of the area under the curves. N=6 independent injections accounting 2 biological replicates representing a total of n=75 cells (WT/HA_Tr.), n=55 cells (KO/HA_Tr.) and n=75 cells (KO/CDK4_Tr.). RM 1 way ANOVA and Tukey’s multiple comparisons test.

To extend these results, we then analyzed calcium flux and signaling between two important intracellular calcium storage organelles, namely, the ER and mitochondria. We used two different inducers of ER calcium release, *i.e.*, thapsigargin (TG) and histamine, and observed mitochondrial and cytosolic calcium kinetics. Both thapsigargin and histamine induced clear accumulation of mitochondrial calcium in CDK4-WT cells but not in CDK4-KO cells (Fig 4E-H and Sup. Fig 4C-D). CDK4-WT and CDK4-KO cells exhibited equivalent ER calcium release kinetics when the cytosolic calcium level was measured in response to the inducers, suggesting that the observed attenuation of mitochondrial calcium uptake was not caused by deficient ER calcium release (Fig 4I-L and Sup. Fig 4C-D). On the other hand, most of the main ER-MT-associated calcium channels except ITPR2 showed either no change in protein expression (SERCA1 and MCU) or increased protein expression (ITPR1, ITPR3 and VDAC1) in CDK4-KO cells (Fig 4M-N). Finally, re-expression of CDK4 in CKD4-KO cells restored the induction of mitochondrial calcium release in response to thapsigargin (Fig 4O-P) (Sup. Fig 4G). These results suggest that CDK4 is required for ER-MT calcium signaling in TNBC cells.

### CDK4 participates in the establishment of MERCs

ER-MT calcium transfer involves membrane contact sites between the ER membrane and the outer mitochondrial membrane (OMM), known as MERCs^17^. These MERCs are also required for the initiation of mitochondrial fission, during which the ER physically wraps mitochondria ^26,27^. Since we observed both reduced ER-MT calcium signaling and decreased mitochondrial fission in CDK4-KO cells (Fig. 4), we next assessed whether the structure of MERCs is modified in CDK4-KO cells using complementary technical approaches, such as TEM, proximity ligation assay (PLA) and coimmunofluorescence staining. TEM analysis showed a decrease in the number of MERCs per mitochondrion in CDK4-KO TNBC cells (Fig 5A-B), which was correlated with an overall decrease in the percentage of MAMs (Fig 5C). Moreover, both the mean and minimum ER-mitochondria distances of the MERCs were increased in CDK4-KO cells compared to CDK4-WT cells (Fig 5D and Sup. Fig 5A), indicating looser contacts between mitochondria and the ER. Accordingly, the xenograft tumors also exhibited an overall decrease in the number of MERCs per mitochondrion, according to the electron micrographs (Sup. Fig 5B). The decreased mitochondria-ER association was further confirmed by tracking mitochondria and the ER and analyzing the subsequent fluorescence colocalization (Sup. Fig 5C). Finally, the PLA results indicated a decrease in close appositions (<50 nm) between ITPR1 (an ER-resident protein) and VDAC1 (an OMM-resident protein) (Fig 5E). Exogenous reintroduction of CDK4 in CDK4-KO cells restored the MERC interface, as shown by the increased number of ITPR1-VDAC1 puncta in the PLA (Fig 5E).

**Figure 5:**
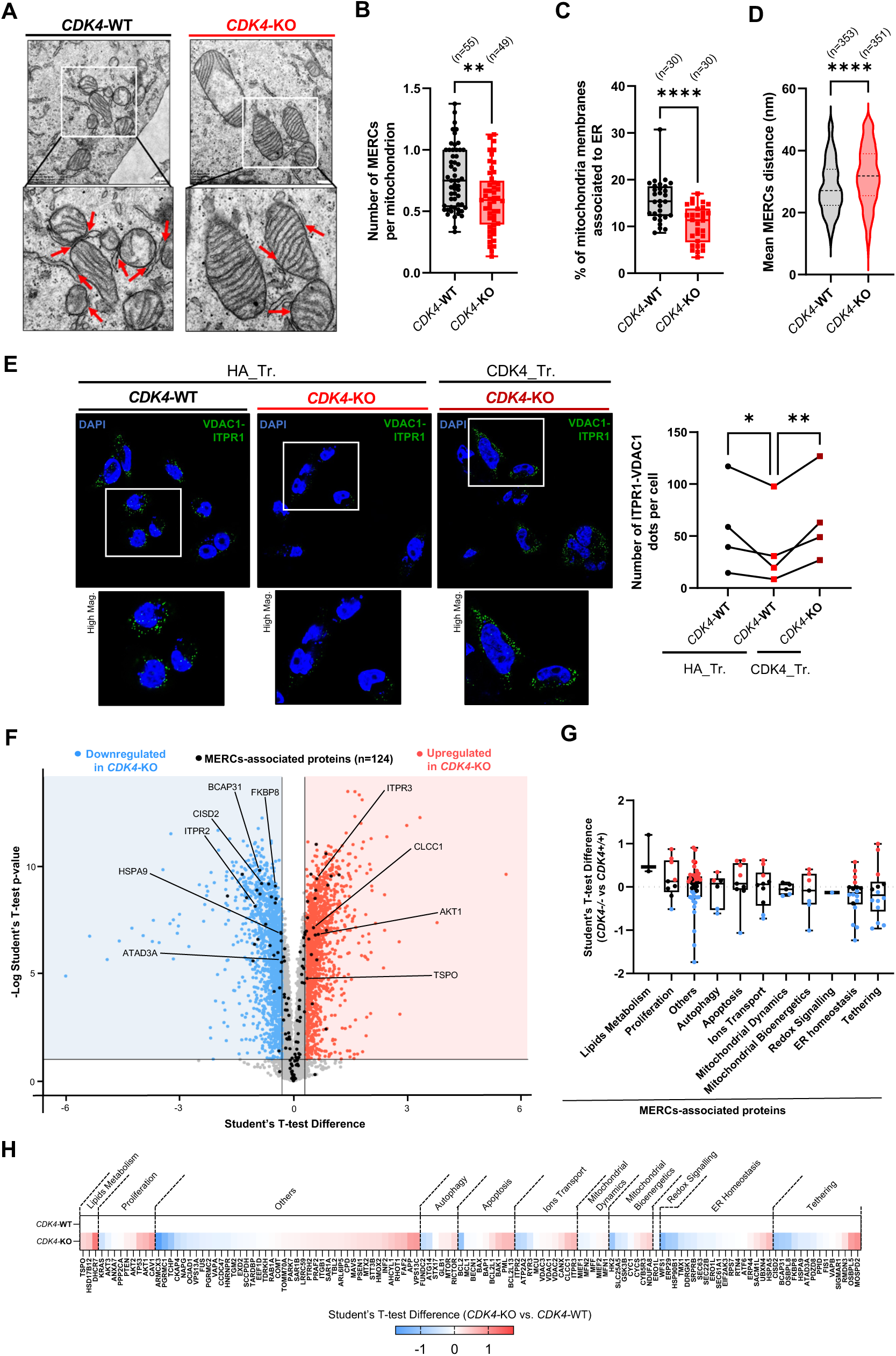
CDK4 enhances Mitochondria-ER Contacts in TNBC. **(A).** Representative electron micrographs of CDK4-WT and CDK4-KO TNBC cells. Scale bars: 500 nm. Red arrows indicate Mitochondria-ER Contacts (MERCs). **(B).** Quantification of the number of MERCs per mitochondrion of CDK4-WT and CDK4-KO TNBC cells. N=3 independent biological replicates representing a total of n (WT) = 55 cells and n (KO) = 49 cells. Mann-Whitney Test. **(C).** Percentage of mitochondria membranes associated to ER. N=3 independent biological replicates representing a total of n (WT) = 55 cells and n (KO) = 49 cells. Mann-Whitney Test. **(D).** Mean ER-mitochondrion distance in analyzed MERCs. N=3 independent biological replicates representing n (WT) = 353 MERCs from n=55 cells and n (KO) = 351 MERCs from n=49 cells. Mann-Whitney Test. **(E).** Proximity Ligation Assay (PLA) using VDAC1 and ITPR1 antibodies in CDK4-WT, CDK4-KO TNBC cells transfected with empty plasmid HA (HA_Tr.) and CDK4-KO TNBC cells expressing endogenous CDK4 (CDK4_Tr.). Representative pictures and associated quantification of ITPR1-VDAC1 dots per cell. N=4 independent biological replicates. 2-way ANOVA and Sidák’s multiple comparison tests. **(F).** Volcano plots of proteins differentially regulated in CDK4-WT and CDK4-KO TNBC cells, highlighting proteins significantly down regulated (blue) and up-regulated (red) in CDK4-KO compared to CDK4-WT TNBC cells. N=5 biological replicates. In black are indicated 124 MERCs-associated proteins found in the proteomic analysis. **(G).** Histogram recapitulating the subclustered classes of the 124 MERCs proteins, from the most upregulated one (left) to the most down-regulated one (right) according to the median of Student’s T-test difference. **(H).** Heatmap of subclustered classes of the 124 MERCs proteins, from the most upregulated one (left) to the most down-regulated one (right) according to the median of Student’s T-test difference.

To better understand how CDK4 may impact the composition of MERCs, we performed a large-scale proteomic analysis in both CDK4-WT and CDK4-KO cells. Based on a meta-analysis of various proteomic profiles of MERC fractions isolated from human cells^28–32^, we first established a list of 156 core MERC proteins (Sup. Table 1). In our data, we detected 124 of these 156 MERC proteins (79%) (Fig 5F). Of these 124 detected proteins, 86 (69%) were differentially regulated upon CDK4 deletion, and most of these differentially regulated proteins were downregulated (Fig 5F). To more precisely delineate this mechanism of CDK4 regulation, the MERC proteins were clustered according to their functions (Sup. Table 1). Strikingly, we found that the MERC tether protein category showed the greatest downregulation in CDK4-KO cells compared to -WT cells (Fig 5G-H). In contrast, lipid metabolism and proliferation-associated proteins were most commonly upregulated in CDK4-KO cells (Fig 5G-H), probably stemming from compensatory mechanisms activated to allow the cells to rely on a smaller number of MERCs but with a high abundance of pro-proliferative and lipid/calcium signaling proteins.

### CDK4 regulates PKA signaling at MERCs

Since CDK4 is a serine-threonine kinase, we hypothesized that it exerts its effects *via* the phosphorylation of new targets implicated in the control of MERCs. We therefore performed PamGene phosphoproteomic analysis coupled to Integrative Inferred Kinase Activity (INKA) analysis^33^. The results of these two independent and complementary analyses converged to identify PKA alpha as the kinase with the most downregulated kinase activity in CDK4-KO cells compared to CDK4-WT cells (Fig 6A and Sup. Fig 6A). Phospho-PKA substrates were consistently downregulated in CDK4-KO cells (Fig 6B). Reintroduction of CDK4 into CDK4-KO cells was sufficient to restore PKA activity (Fig 6C), which proved a direct effect of CDK4. We further assessed whether the kinase activity of CDK4 was needed for maintaining PKA activity. To this end, we used a kinase-dead CDK4 mutant (K35M) and a non-inhibitable CDK4 mutant (R24C; to rescue CDK4 activity in KO cells) and revealed that CDK4 K35M could not reverse the decreases in the expression of phospho-PKA substrates, in contrast to CDK4 R24C (Fig 6C), proving that the kinase activity of CDK4 is required for PKA activity. Moreover, the phosphorylation of ITPR1 at S1756, which is a known PKA target site and an important potentiating-phosphorylation of ITPR-evoked calcium release ^34^ was consistently decreased in CDK4-KO cells (Fig 6D-E).

**Figure 6:**
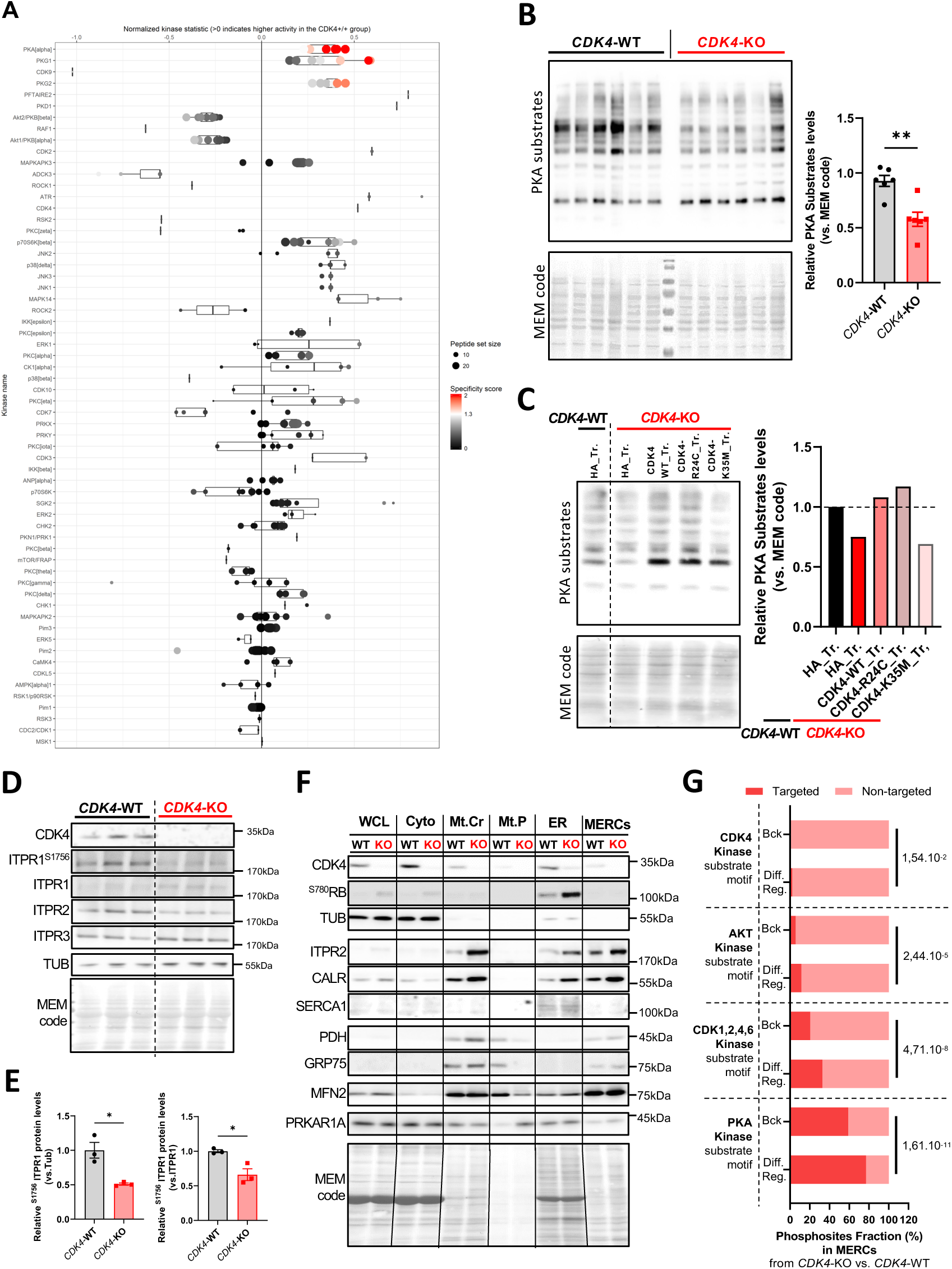
CDK4 regulates PKA signaling at ER-MT. **(A).** Kinome profiling/PAMgene analysis CDK4-WT and CDK4-KO TNBC cells. Comparison CDK4-WT vs. CDK4-KO TNBC cells. **(B).** Immunoblots of Phospho-PKA substrates and MEM code in CDK4-WT and CDK4-KO TNBC cells. Relative quantity of Phospho-PKA substrates normalized to MEM code. N=6 independent biological replicates. **(C).** Immunoblots of Phospho-PKA substrates and MEM code of CDK4 WT and CDK4 KO TNBC cells. Relative quantity of Phospho-PKA substrates relative to MEM code. Representative of N=3 independent biological replicates. **(D-E.)** Immunoblots and relative protein levels of CDK4, ^S1756^ITPR1, ITPR1, ITPR2, ITPR3 and Tubulin (TUB) and MEM code of CDK4-WT and—KO TNBC cells. N=3 independent biological replicates. Unpaired T-tests. **(F).** Immunoblots of CDK4, ^S780RB^, Tubulin (TUB), ITPR2, CALR, SERCA1, PDH, GRP75, MFN2, PRKAR1A and MEM code in subcellular fractions of CDK4-WT and—KO TNBC cells. WCL=Whole cell lysate. Cyto=Cytosolic fraction. Mt.Cr=Crude mitochondria fractions. Mt.P=Pure mitochondria fractions. ER=Endoplasmic reticulum fraction. MERCs=Mitochondria-ER contacts fractions. **(G).** Enrichment analysis on phospho-peptides found in MERCs fraction of CDK4-WT and—KO TNBC cells. Phosphosites displaying CDK4 kinase motifs, AKT kinase motifs, CDK1,2,4,6-kinase motifs or PKA kinase motifs are represented. Bck=Background. Diff.Ref.=Differentially regulated. N=3 replicates. Benj. Hoch. FDR value is displaying for each substrate motif category.

PKA is a tetrameric holoenzyme composed of two regulatory and two catalytic subunits and is inactive in this state ^35^. The PKA enzymatic complex is organized by AKAP anchoring proteins, which facilitate its interactions, including those of the regulatory subunits and its substrates. AKAPs are also important for the specific intracellular localization of PKA. Upon the binding of cAMP to the PKA regulatory subunits, the catalytic subunits are released from the complex, and PKA is activated. To identify PKA substrates and elucidate the localization of PKA at MERCs, we performed proteomic and phosphoproteomic analyses in isolated crude/pure mitochondria, ER, and MERC fractions. The purity of the fractions was validated using specific markers for the ER, mitochondria and MERC fractions (Fig 6F). Of note, CDK4 was detected in the cytosolic, mitochondrial, ER and MERC fractions (Fig 6F). A total of 3659 proteins were enriched in the MERC fraction, and identified 101 core MERC proteins in our dataset. Notably, based on the list of 154 MCC core proteins established with previous proteomics data (Sup. Table 1), we detected 101 of these MERC proteins in the MERC fraction, with 38 (25%) detected only in the MERC fraction and 63 (41%) enriched in the MERC fraction compared to the whole-cell lysate (37.6%) (Sup. Fig 6B). Proteomic analysis also revealed that most of these 101 core MERC proteins were upregulated in CDK4-KO cells (Sup. Fig 6C), suggesting a compensatory effect on the decrease in the number of MERCs in these cells. Interestingly, the MERC fraction contained at least two regulatory subunits (PRKAR1A, PRKAR2A), two catalytic subunits (PRKACA, PRKACB) and four AKAPs (AKAP1, 2, 5 and 10) (Sup. Fig 6C). We confirmed the localization of the PKA regulatory subunit PRKAR1A in MERCs (Fig 6F) and the presence of PKA phosphorylation substrates, which were deregulated in the MERC fraction of CDK4-KO cells (Sup. Fig 6D). Furthermore, phosphoproteomic enrichment analysis demonstrated that phosphopeptides with PKA motifs were significantly deregulated in the MERCs of CDK4-KO cells, and this deregulation was correlated with the deregulation of phosphopeptides containing a CDK4 phosphorylation motif (Fig 6G and Sup. Fig 6F). In addition to identifying PKA-targeted phosphosites, we identified 59 differentially regulated phosphosites in 49 MERC-associated proteins (Sup. Fig 6E). Among these different phosphorylation targets, many calcium-related channels involved in ER-MT calcium transfer, including ITPR1, ITPR2, ITPR3, VDAC1 and VDAC2, were downregulated in CDK4-KO cells (Sup. Fig 6E).

### CDK4 promotes mitochondrial fitness and metabolic flexibility in TNBC cells

MERCs regulate mitochondrial bioenergetics through calcium fluxes, which are determinants of the activity of many matrix dehydrogenases, such as pyruvate dehydrogenase (PDH), α-ketoglutarate dehydrogenase (αKGDH) or isocitrate dehydrogenase (IDH)^17,18,36^. As CDK4-KO cells display altered MERC tethering and attenuated ER-MT calcium signaling, we evaluated the functionality of their mitochondria. We first observed that the giant mitochondria in CDK4-KO cells had an altered cristae structure (Fig 7A), with an increased number of cristae per mitochondrion (Fig 7B). This increase correlated with the overall decrease in the length of cristae normalized to the mitochondrial area (Fig 7B). In addition to investigating these markers of crista remodeling, we evaluated the global mitochondrial membrane potential (MMP) using the ratio of MitoTracker Red (MMP-dependent) to MitoTracker Green (MMP-independent) fluorescence (Fig 7C). Remarkably, CDK4 depletion resulted in a reduction in this ratio, indicating a decreased MMP in CDK4-KO cells compared to CDK4-WT cells (Fig 7C).

**Figure 7:**
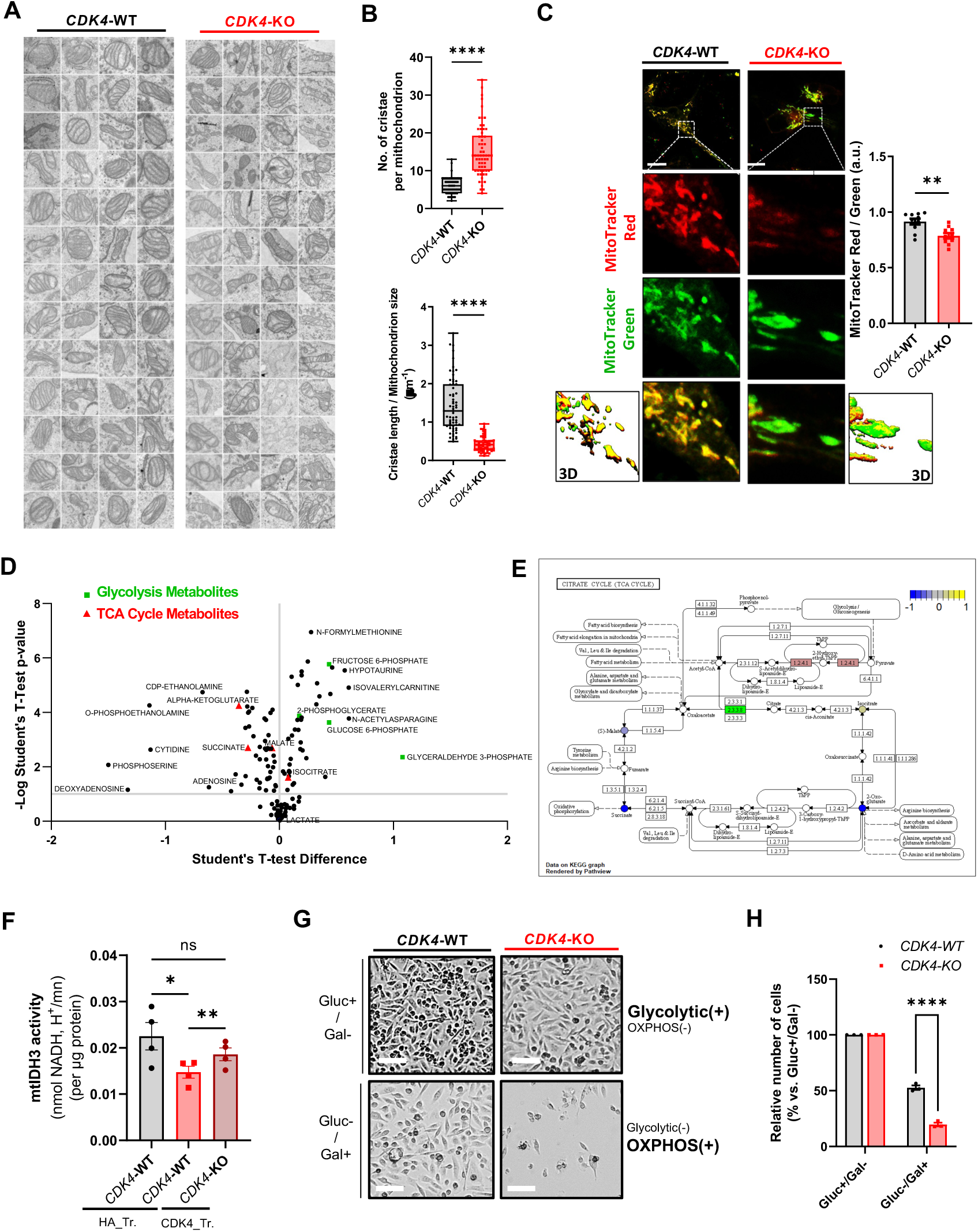
CDK4 promotes mitochondrial fitness and metabolic flexibility through balanced MT calcium signaling. **(A).** Representative electron micrographs of mitochondria cristae from CDK4-WT and CDK4-KO TNBC cells. N=3 independent biological replicates representing n=52 mitochondria. **(B).** Quantification of the number of cristae per mitochondria and cristae length/mitochondria area from CDK4-WT and CDK4-KO TNBC cells. N=3 independent biological replicates representing n=52 mitochondria. Mann-Whitney Test. **(C).** Representative micrographs of CDK4-WT and CDK4-KO TNBC cells stained with MitoTracker Red (MMP-dependent) and MitoTracker Green (MMP-independent). Quantification of MitoTracker Red on MitoTracker Green fluorescence intensities in CDK4-WT and CDK4-KO TNBC cells. n=12 cells. Scale bars: 10 μm. **(D).** Metabolomics analysis from multi-pathway analysis from CDK4-WT and CDK4-KO TNBC cells. Metabolism from glycolysis pathways (blue), TCA cycle (red), and glutamine metabolism (green) are highlighted. N=5 biological replicates. **(E).** TCA cycle scheme with annotated metabolites found in metabolomics analysis. The relative quantity of TCA metabolites is represented. Main calcium-dependent enzymes are represented in bold. N=5 biological replicates. Multiple Student’s t-test **(F).** Mitochondrial Isocitrate-dehydrogenase 3 (mtIDH3) activity normalized by the quantity of proteins and measured in CDK4-WT, CDK4-KO TNBC cells transfected with empty plasmid HA (HA_Tr.) and CDK4-KO TNBC cells expressing endogenous CDK4 (CDK4_Tr.). N=4 biological replicates.). RM 1 way ANOVA and Tukey’s multiple comparisons test. **(G)**. Representative pictures of CDK4-WT and CDK4-KO TNBC cells 3 days after continuous culture with glucose (+ + + = 20 mM)/No Galactose (-) or galactose (+ + + = 10 mM)/no glucose (-) media.

To assess the changes in the main metabolites in TNBC cells upon CDK4 deletion, we performed targeted mass spectrometry-based analysis of 200 polar and moderately metabolites in CDK4-WT and CDK4-KO cell extracts. Interestingly, CDK4-KO cells showed enrichment of glycolytic metabolites compared to CDK4-WT cells (Fig 7D). Furthermore, the levels of four tricarboxylic acid (TCA) cycle metabolites were significantly altered in CDK4-KO cells (Fig 7D). Among them, isocitrate level was highly increased, in contrast to α-ketoglutarate, succinate and malate, which were depleted in CDK4-KO cells (Fig 7D-E), suggesting a blockade in the conversion of isocitrate into α-ketoglutarate *via* IDH. As calcium regulates the activity of mitochondrial-dehydrogenase enzymes ^36^, we assessed the activity of mitochondrial isocitrate dehydrogenase 3 (IDH3). The activity of mitochondrial IDH3 was consistently decreased in CDK4-KO cells (Fig 7F). Importantly, exogenous reintroduction of CDK4 partially rescued this attenuated activity in CDK4-KO cells (Fig 7F). Finally, we elucidated the differential response of CDK4-KO mice to mitochondrial metabolic challenge. Replacement of glucose with galactose, resulting in inhibition of glycolysis and forcing reliance on oxidative phosphorylation (OXPHOS) for ATP generation, drastically reduced the viability of CDK4-KO cells (Fig. 7G-H). Taken together, these data indicate that CDK4-KO cells, in addition to their ability to resist apoptotic stimuli, exhibit mitochondrial metabolic vulnerabilities that could be mediated by reduced MERC establishment and impaired ER-MT calcium signaling.

## Discussion

In general, when cells divide, they become sensitive to most chemotherapeutic agents, whereas resting cells are resistant. Paradoxically, although CDK4/6i have cytostatic effects, previous studies proved that TNBC and ovarian cancer cells treated with doxorubicin, paclitaxel, or carboplatin were resistant to cytotoxic effects and the associated cell death in response to CDK4/6 inhibition ^37–39^. In fact, inhibition of CDK4/6 increases the expression of *BCL2L1* (encoding Bcl-xL) and *MCL1*, which encode two antiapoptotic members of the Bcl-2 protein family, ultimately resulting in apoptosis evasion ^40^. CDK4/6 inhibition promotes the survival of kidney cells upon cisplatin or etoposide treatment ^41–43^ and of normal intestinal cells upon radiation-induced injury ^44^. Furthermore, the use of CDK4/6i increases the tolerability of chemotherapy in lung cancer patients, as observed by the diminished hematological toxicity and myelopreservation in multiple hematopoietic lineages ^45^. Collectively, these data suggest that CDK4/6 inhibition protects cells against chemotoxicity. Furthermore, recent data from the Malumbres laboratory showed that pretreatment of cancer cells with CDK4/6i rendered the cells resistant to chemotherapeutic drugs, whereas treatment after chemotherapy resulted in synergistic effects ^46^.

In our study, we reproduced the resistance to CDK4i using an *in vivo* xenograft model established with CDK4-KO TNBC cells. The dynamics of the acquisition of resistance were indeed very similar (Fig 1I in our manuscript and Fig 7E in ^47^), strongly suggesting that CDK4 plays a prominent role in the development and therapeutic resistance in this particular cancer. Furthermore, we proved that the formation of CDK4-KO tumors in xenografted mice does not result from CDK4 re-expression that could be secondary to acquired mutations (Fig 1K). It is, however, puzzling why there is a latency period of approximately 21 days, in which, similar to treated cells ^47^ CDK4-KO TNBC cells do not grow in grafted mice. This adaptation period may depend on the specific metabolic needs of these cells that are required for the development of the tumor microenvironment.

Another important finding of our study was the differences in mitochondrial dynamics and function in CDK4-KO cells, which add another layer of complexity to the understanding of the role of CDK4 in TNBC and possibly in other cancers. Mitochondria are vital organelles responsible for energy production, regulation of apoptosis, and various other cellular functions. Alterations in mitochondrial dynamics and function often have profound implications for cell survival and metabolism ^48,49^. CDK4-KO cells exhibit defects in mitochondrial morphology that are the result of dysregulated mitochondrial dynamics, which can lead to impaired energy production, calcium signaling, and altered apoptotic responses.

The underlying mechanisms explaining the antiapoptotic effects of CDK4 deletion in TNBC include the regulation of mitochondrial biology. The differences in mitochondrial dynamics and function in CDK4-KO cells are linked to their apoptosis resistance. We demonstrated that in response to distinct apoptotic stresses, robust mitochondrial calcium uptake, which was observed in WT cells, was not elicited in CDK4-KO TNBC cells. An increase in intramitochondrial calcium is required for the induction of apoptosis. An elevated calcium level can lead to opening of the mitochondrial permeability transition pore (mPTP), disrupting the MMP, dissipating the proton gradient, and finally allowing the release of cytochrome c, which then participates in the activation of caspase-9, an initiator protein in the intrinsic apoptotic pathway ^50^.

The uptake of calcium into mitochondria is mediated mainly by mitochondria-endoplasmic reticulum contacts. MERCs constitute 10 to 20% of all mitochondrial membranes, highlighting not only their importance for cellular functions but also their dynamic nature, as this percentage varies across species, tissues and cell types ^22^. We demonstrated in this study that CDK4 localizes at MERCs, modulating their tethering and influencing their signaling through PKA misregulation. Indeed, the decreased number of MERCs in CDK4-KO cells is translated into decreased uptake of calcium into mitochondria in response to apoptotic stimuli, thereby protecting cells from death.

We identified the protein kinase PKA as a target of CDK4 at the MERC interface. A recent study suggested that PKA promotes mitochondrial remodeling and ER-MT calcium signaling during the early phase of ER stress ^51^. This enhanced ER-MT calcium signaling thus favors mitochondrial bioenergetics during the adaptive response. PKA signaling is a determinant of MERC formation, as it mediates the phosphorylation of key proteins involved in calcium signaling and mitochondrial fission, such as ITPR1 ^34^ and the pro-fission protein DRP1 ^52^, respectively. AKAP1, which mediates PKA subunit translocation to the OMM, also localizes at MERCs^53^, reinforcing the importance of PKA signaling in this subcellular compartment. Here, we demonstrated an upstream mechanism of PKA regulation through CDK4 kinase activity. Of note, in the analysis of PKA motif phosphosites differentially regulated at MERCs, we identified many other calcium channels, such as VDAC1, and this finding may facilitate the understanding of the biological complexity of mitochondrial PKA signaling^35,54,55^. Furthermore, the regulation of PKA activity at the MERC interface may facilitate the understanding of how and why the activities of mitochondrial and cytosolic PKA have opposite effects on apoptosis, either promoting or attenuating it, respectively.

Mitochondrial dysfunction in CDK4-KO TNBC cells, which was characterized by impaired mitochondrial dynamics and calcium signaling, had other physiological consequences in addition to protection against cell death. CDK4 deletion in these cells also compromised the metabolic flexibility required for the response to metabolic challenges to efficiently drive mitochondrial adaptive metabolism, as observed during cancer progression. Previous reports have provided evidence that OXPHOS and mitochondrial functions constitute metabolic vulnerabilities in metastatic breast cancers with resistance to the CDK4 inhibitor palbociclib ^56^. Similar effects were also observed in pancreatic cancer ^57^, colorectal cancer^58^, and melanoma^59^. In our model, deletion of CDK4 also resulted in metabolic vulnerabilities in TNBC cells, which could not grow in the presence of galactose, indicating defective mitochondrial function. Moreover, using metabolomic analysis coupled to gene expression analysis, we found that the activity of mitochondrial IDH3 was significantly decreased in CDK4-KO cells. Consistent with these results, the levels of both α-ketoglutarate and succinate, which are downstream metabolites derived from isocitrate in the TCA cycle, were decreased in these cells.

In summary, this study underscores a new role for CDK4 in the regulation of cell death through the control of MERCs, directly impacting mitochondrial dynamics and calcium signaling. The identification of this new function could lead to the establishment of a new paradigm in the treatment of metastatic breast cancer and possibly other cancers through consideration of how CDK4i treatment could promote resistance to chemotherapy. Our findings also pave the way for defining new therapeutic strategies based on the metabolic vulnerabilities proven to be present when CDK4 is inhibited.

## Material and methods

### Cell culture, plasmid transfection and probes

MDA-MB-231 TNBC cells (ATCC) were cultured in Roswell Park Memorial Institute (RPMI)-1640 medium with GlutaMAX (Gibco^TM^, Ref: 61870036) supplemented with 10% heat-inactivated FBS (HyClone, Cat. No: SV30160, Lot: RB35956), 10 mM HEPES (Gibco^TM^, Ref: 15630122) and 1 mM pyruvate (Gibco^TM^, Ref: 11360070). Cells were grown under normoxic conditions at 37°C and 5% C0_2_. The method used to generate CDK4-KO cells *via* CRISPR/Cas9 gene editing is described in ^10^. Empty-HA (Ctrl), CDK4-WT, CDK4-R24C, and CDK4-K35M plasmids were provided by Marcos Malumbres (CNIO, Spain). The pFusionRed-ER vector (Evrogen, #FP420) was used for ER tracking in cells. For transfection, cells were plated one day before transfection and were then incubated with a 2:1 ratio (μL:μg) of X-tremeGENE™ HP DNA Transfection Reagent (Roche, Ref: 6366236001) and plasmids for 20 min in Opti-MEM (Gibco^TM^, Ref: 51985034). To track total and active mitochondria, we incubated cells with MitoTracker Green (Invitrogen^TM^, Ref: M7514) and MitoTracker Red (Invitrogen^TM^: Ref: M22425) (200 nM), respectively, in Opti-MEM (Gibco^TM^, Ref: 51985034) for 30 min at 37°C and 5% CO_2_.

### Animals

NOD SCID gamma (NSG) mice from The Jackson Laboratory were bred at the animal facility of the University of Lausanne. 2 x 10^6^ MDA-MB-231 CDK4-WT or -KO cells in 50μL of 1X PBS were injected into the fat pad of the 4^th^ mammary gland of 8 week-old females. Tumor volume was assessed by measuring the length and the width with a caliper. The mice received Cisplatin (0,8mg/kg) by intraperitoneal injection when the tumors reached a mean size volume of 80 mm^3^, and the treatment was repeated every 4 days during 12 days. The experiments were approved by the veterinary office of the State of Vaud (Cantonal License #VD3726, National License #34118), and the conducted methods were in accordance with the animal care guidelines of Swiss laws, following 3R recommendations.

### RT-qPCR and RNA sequencing

Reads were mapped to the mouse genome GRCh38 (Ensembl version 102) using STAR (v. 2.7.0f; options: --outFilterType BySJout –outFilterMultimapNmax 20 –outMultimapperOrder Random –alignSJoverhangMin 8 –alignSJDBoverhangMin 1 – outFilterMismatchNmax 999 –alignIntronMin 20 –alignIntronMax 1000000 – alignMatesGapMax 1000000). Read counts in gene loci were evaluated with HTSeq-count (v. 0.13.5). Differential expression analysis was performed in R (v 4.1.1) using the DESqe2 package with p value adjustments for multiple comparisons (Benjamini‒Hochberg procedure). Principal component analysis (PCA) was performed in R using variance-stabilized expression data for the 500 genes with the greatest variability. GSEA was performed in R using MsigDB and the fgsa Bioconductor package. Kyoto Encyclopedia of Genes and Genomes (KEGG) and Gene Ontology (GO) enrichment analyses were performed in R using the ReactomePA Bioconductor package. For RT-qPCR, RNA was isolated by the Trizol-Chloroforme method extraction and subsequent isopropanol-based precipitation. Reverse transcription was performed with 400ng of RNA using SuperScript™ II Reverse Transcriptase (ThermoFisher). 5 ng of cDNA was amplified with specific primers and Maxima™ SYBR green/ROX QPCR master mix (2X) (Thermofisher). The primers sequences are listed in Supplementary Table S2.

### Proteomics and Phospho-Proteomics

Total cells were lysed in RIPA buffer, 2% SDS, 10mM DTT. Tryptic Digestion was done following the SP3 method^60^ using magnetic Sera-Mag Speedbeads (Cytiva 45152105050250, 50 mg/mL). After heating for 10 min at 75°C, cysteines were alkylated with 32mM (final) iodoacetamide for 45 min at RT in the dark. Beads were added at a ratio 10:1 (w:w) to samples, and proteins were precipitated on beads with ethanol (final concentration: 60 %). After 3 washes with 80% ethanol, on-beads digestion was performed in 40µl of 50mM Hepes pH 8.3 with 2µg of trypsin (Promega #V5073). Supernatants were harvested for TMT labelling. Supernatants from SP3 digestion containing each 100µg peptides in 50 µL Hepes pH 8.3 buffer were mixed each with 0.4 mg of TMT 10-plex reagent (Thermo Fisher Scientific product nr 90111) dissolved in anhydrous acetonitrile and incubated at RT for 90 min. Excess reagent was quenched by adding 7µl of hydroxylamine 5% (v:v) and incubating 15 min at RT. After mixing and evaporation of excess acetonitrile, samples were acidified and desalted on SepPak C18 cartridges. Peptides were eluted in 50% acetonitrile and dried. Aliquots (1/10) of samples were analysed directly by LC-MS to determine total proteome composition. Phosphopeptide enrichment by IMAC was performed on the remaining of TMT-labelled material (a16pprox. 0.4-0.6 mg) with the High-Select™ Fe-NTA Phosphopeptide Enrichment Kit (Thermo Fisher Scientific, Prod. Number A32992) according to kit instructions. IMAC eluates were dried and resuspended for analysis in 2% MeCN, 0.05% TFA. Data-dependent LC-MS/MS analyses of samples were carried out on a Fusion Tribrid Orbitrap mass spectrometer (Thermo Fisher Scientific) connected through a nano-electrospray ion source to an Ultimate 3000 RSLCnano HPLC system (Dionex), *via* a FAIMS interface. Peptides were separated on a reversed-phase custom packed 45 cm C18 column (75μm ID, 100Å, Reprosil Pur 1.9µm particles, Dr. Maisch, Germany) with a 4-90% acetonitrile gradient in 0.1% formic acid (140 min). Data files were analysed with MaxQuant 1.6.14.0 incorporating the Andromeda search engine^61^. Cysteine carbamidomethylation and TMT labelling (peptide N-termini and lysine side chains) were selected as fixed modification while methionine oxidation and protein N-terminal acetylation were specified as variable modifications. Both peptide and protein identifications were filtered at 1% FDR relative to hits against a decoy database built by reversing protein sequences. For TMT analysis, the raw reporter ion intensities generated by MaxQuant (with a mass tolerance of 0.003 Da) and summed for each protein group were used in all following steps to derive quantitation. All subsequent analyses were done with the Perseus software package (version 1.6.15.0). Contaminant proteins were removed. After imputation of missing values (based on normal distribution using Perseus default parameters), t-tests were carried out among all conditions, with permutation-based FDR correction for multiple testing (Q-value threshold <0.01). The difference of means obtained from the tests were used for 1D enrichment analysis on associated GO/KEGG annotations as described^62^. The enrichment analysis was also FDR-filtered (Benjamini-Hochberg, Q-val<0.02). The mass spectrometry proteomics data including raw output tables have been deposited to the ProteomeXchange Consortium *via* the PRIDE partner repository ^63^ with the dataset identifiers PXD046326, PXD046327 and PXD046353.

### Kinome profiling (PamGene)

For kinome analysis, serine/threonine kinase (STK) microarrays were purchased from PamGene International BV. Each array contained 140 phosphorylatable peptides as well as 4 control peptides. Sample incubation, detection, and analysis were performed according to the manufacturer’s instructions in a PamStation 12 instrument. In brief, extracts from BMDMs or human visceral adipose tissue (VAT) were prepared using M-PER mammalian extraction buffer (Thermo Scientific) containing Halt phosphatase inhibitor cocktail (1:50; Thermo Scientific) and Halt protease inhibitor cocktail, EDTA-free (1:50, Thermo Scientific) for 20 min on a rotating wheel at 4°C. The lysates were then centrifuged at 13,000 rpm for 20 min to remove all debris. The supernatants were aliquoted, snap-frozen in liquid nitrogen, and stored at -80°C until further processing. Prior to incubation with the kinase reaction mix, the arrays were blocked with 2% BSA for 30 cycles and washed three times with PK assay buffer. Kinase reactions were performed over a 1-h period with 5 µg of total extract and 400 µM ATP at 30°C. Phosphorylated peptides were detected with a -FITC-conjugated anti-rabbit secondary antibody that recognizes a pool of anti-phosphoserine/threonine antibodies. The PamStation 12 instrument contains a 12-bit CCD camera suitable for imaging of FITC-labeled arrays. The images of the phosphorylated arrays were used for quantification with BioNavigator software (PamGene International BV). The generated heatmaps and BioNavigator score plots are further explained in the results section and figure legends.

### Electron microscopy

Mouse tissue was cut into 1 mm^3^ pieces and fixed with 2.5% glutaraldehyde solution (EMS, Hatfield, PA, US) in phosphate buffer (PB; 0.1 M, pH 7.4) (Sigma, St. Louis, MO, US) for 1h at room temperature (RT). Then, the samples were rinsed 3 times for 5 min each in PB buffer and postfixed with a fresh mixture of 1% osmium tetroxide (EMS, Hatfield, PA, US) and 1.5% potassium ferrocyanide (Sigma, St. Louis, MO, US) in PB buffer for 1h at RT. The samples were then washed three times in distilled water and dehydrated in acetone solution (Sigma, St. Louis, MO, US) at graded concentrations (30%, 40 min; 70%, 40 min; 100%, 1 h; 100%, 2 h). This step was followed by infiltration in Epon resin (Sigma, St. Louis, MO, US) at graded concentrations (Epon 1/3 acetone, -2 h; Epon 3/1 acetone, -2 h, Epon 1/1, -4 h; Epon 1/1, -12 h) and polymerization for 48 h at 60°C in an oven. Thin sections of 50 nm were sliced on a Leica Ultracut microtome (Leica Mikrosysteme GmbH, Vienna, Austria) and picked up on a copper grid (2×1 mm; EMS, Hatfield, PA, US) coated with a polyetherimide (PEI) film (Sigma, St. Louis, MO, US). Sections were sequentially poststained with 2% uranyl acetate (Sigma, St. Louis, MO, US) in H_2_O for 10 min, rinsed several times with H_2_O followed by Reynolds lead citrate for 10 min and rinsed several times with H_2_O. Large montages were acquired with a Philips CM100 transmission electron microscope (Thermo Fisher Scientific, Hillsboro, US) at an acceleration voltage of 80 kV with a TVIPS TemCam-F416 digital camera (TVIPS GmbH, Gauting, Germany), and alignment was performed using the Blendmont command-line program in IMOD software (Kremer, Mastronarde, et McIntosh 1996).

### MERC isolation

A total of 100×10^6^ *CDK4*-WT and *CDK4*-KO MDA-MB-231 TNBC cells were plated for isolation of subcellular fractions as described in^64^.

### Calcium signaling

For cytosolic calcium measurement, cells were loaded with 1 mL of Krebs solution with calcium (mM: NaCl 135.5, MgCl_2_ 1.2, KCl 5.9, glucose 11.5, HEPES 11.5, CaCl_2_ 1.8; final pH, 7.3) containing 5 μM of the cytosolic Ca^2+^ indicator Fluo-4 AM (Invitrogen, Switzerland) solubilized in Krebs solution (in mM: NaCl 135.5, MgCl_2_ 1.2, KCl 5.9, glucose 11.5, HEPES 11.5, CaCl_2_ 1.8; final pH, 7.3) by incubation for 20 min. The cells were then rinsed twice with calcium-free Krebs solution (in mM: NaCl 135.5, MgCl_2_ 1.2, KCl 5.9, glucose 11.5, HEPES 11.5, 0.2 Na-EGTA; final pH, 7.3). For mitochondrial calcium measurement, cells were incubated with 1 mL of Krebs solution with calcium containing 1 μM of the fluorescent mitochondrial indicator Rhod-2 AM (Invitrogen, Switzerland) for 1 h at RT. The cells were then washed twice with calcium-free Krebs solution. Both Fluo-4 and Rhod-2 AM fluorescence were monitored using a Zeiss LSM 780 Live confocal microscope system with a 40× oil immersion lens. For Fluo-4 AM, the excitation wavelength was set at 488 nm, and the emitted fluorescence was collected through a bandpass filter (495-525 nm). For Rhod-2 AM, the excitation wavelength was set at 532 nm, and the emitted fluorescence was collected through a bandpass filter (540–625 nm). Thapsigargin (2 μM) or histamine (50 μM) was independently added 60 sec after starting the acquisition of the region of interest (ROI) for each individual cell. Changes in Fluo-4 and Rhod-2 fluorescence were calculated by reporting the peak fluorescence intensity with respect to the baseline fluorescence intensity (as the corresponding relative cytosolic or mitochondrial calcium concentration), equivalent to the mean intensity during the first 60 sec of the acquisition. The mean of one biological replicate included data from 2 to 4 technical replicates with injection (accounting for 20 to 50 analyzed cells).

### Immunoblotting, immunofluorescence staining and proximity ligation assay

For immunoblot analysis, cells were lysed using M-PER™ Mammalian Protein Extraction Reagent (Thermo Scientific, Ref: 78501) complemented with 1X protease and phosphatase inhibitor cocktails (Thermo Scientific, Ref: 78429 and Ref: 1861277). After protein quantification using the Bradford assay, 15-20 μg of protein was loaded onto a gel, resolved by SDS‒PAGE and then transferred to a nitrocellulose membrane (Amersham Protran, Ref: 10600002). Membranes were blocked with 1X TBS -containing 0.05% Tween/3% BSA for 1 h and incubated at 4°C with primary antibodies overnight. The next day, the membranes were washed 3 times with 1X TBS -containing 0.05% Tween and further incubated with a secondary antibody diluted in 1X TBS - containing 0.05% Tween/5% skim milk for 1 h at RT. Membranes were then washed 3 times with 1X TBS -containing 0.05% Tween, and bands were detected using an ECL kit corresponding to the antibodies used (Amersham, Ref: RPN2235 and Advansta, Ref: K-12045-D20).

For immunofluorescence staining of *in vivo* samples, xenograft tumors were harvested, washed with 1X PBS, and fixed for 16 h in 4% paraformaldehyde at +4°C. The xenograft tumors were then washed with 1X PBS and embedded in paraffin. Tumors were sectioned on a Microm HM325 microtome at a thickness of 4 µm. The sections were rehydrated, and antigen retrieval was performed in citrate buffer (pH 6). Following blocking with 5% normal goat serum (in 1X PBS) for 1 h, the cells were incubated overnight at 4°C with the following primary antibodies: anti-mouse cleaved Caspase-3 (Cell Signaling, 9661S; 1:100 dilution) and anti-rat Ki67 (eBioscience 41-5698-80; 1:100 dilution). The following day, the sections were washed in 1X PBS before incubation with Alexa Fluor 488-conjugated goat anti-mouse (1:500) and Alexa Fluor 568-conjugated goat anti-rat (1:500) secondary antibodies for 30 min at RT. The sections were counterstained with DAPI (Invitrogen).

For the PLA, Duolink® reagents were used (Merck, Ref: DUO92014, DUO92002 DUO92004). Cells were washed with 1X PBS and fixed for 10 min in 10% paraformaldehyde. An equal volume of 1 M glycine was added, and the cells were then washed with 100 mM glycine for 15 min. The cells were permeabilized with 0.1% Triton-X-100 and incubated overnight at 4°C with primary antibodies. Cells were washed with 1X PBS containing 0.3% Tween and incubated with PLA probes. Ligation and polymerization were performed according to the manufacturer’s recommendations (Merck, Duolink® procedure). All primary antibodies and dilutions used are listed in Supplementary Table S3.

### Multiple pathway targeted metabolite analysis

Cells were pre-extracted and homogenized by the addition of 1000 µL of MeOH:H_2_O (4:1), in the Cryolys Precellys 24 sample Homogenizer (2 x 20 sec at 10000 rpm, Bertin Technologies, Rockville, MD, US) with ceramic beads. The bead beater was air-cooled down at a flow rate of 110 L/min at 6 bar. Homogenized extracts were centrifuged for 15 min at 4000 g at 4°C (Hermle, Gosheim, Germany). The resulting supernatant was collected and evaporated to dryness in a vacuum concentrator (LabConco, Missouri, US). Dried sample extracts were resuspended in MeOH:H_2_O (4:1, v/v) according to the total protein content evaluated using BCA Protein Assay Kit (Thermo Scientific, Masschusetts, US) Cell extracts were then analyzed by Hydrophilic Interaction Liquid Chromatography coupled to tandem mass spectrometry (HILIC - MS/MS) as previously described by Gallart-Ayala and van der Velpen^65,66^. In brief, to maximize the metabolome coverage, the data were acquired in positive and in negative ionization modes using a 6495 triple quadrupole system (QqQ) interfaced with 1290 UHPLC system (Agilent Technologies).

### IDH activity assay

The IDH Activity Assay Kit (Signa-Aldrich, Ref:MAK062) was used to assess of mitochondrial IDH3 activity. Two days before the assay, one million cells per dish were plated in 10 cm dishes. On the day of the assay, the cells were harvested by scraping with 500 µL of IDH Assay Buffer, and the mixture was centrifuged for 10 min at 13000 rpm and 4°C. Ten microliters of the supernatant was then used to evaluate IDH3 activity according to the manufacturer’s recommendations. Only the cofactor NAD+, preferentially used by mitochondrial IDH3, was added to the reaction buffer mix to distinguish mitochondrial IDH3 activity from the activity of cytosolic and peroxisomal IDH1 and IDH2.

### Data representation and statistical analysis

In bar plots, individual values are presented as the mean ± SEM of N independent biological replicates for *in vitro* experiments or of N mice for *in vivo* experiments. In whisker and violin plots, n indicates the number of mitochondria, MERCs, or cells, as specified in the figure legend. The Shapiro‒Wilk normality test was applied to raw data before proceeding to any analysis. Further statistical analyses and tests are indicated in the figure legends. For comparisons between two groups of non-normally distributed data, the nonparametric Mann‒Whitney test was performed. For comparisons between two groups of normally distributed data, the following two-tailed parametric tests were performed: Student’s t test (equal variance) and Welch’s t test (nonequal variance). For comparisons among more than two groups, repeated measures (RM) (for paired data) or ordinary (for unpaired data) one-way analysis of variance (ANOVA) was used, and Tukey’s multiple comparisons test (with a single pooled variance) was subsequently performed. For data from *in vivo* experiments, two-sided Grubbs’ test was performed to find outliers, which were removed from further analysis if the p value was less than 0.05. All statistical analyses were performed using GraphPad Prism 9.1.0 software (ns: nonsignificant; * p < 0.05; ** p < 0.01; *** p < 0.001).

## Author contributions

D.V.Z. and L.F. conceived and designed the study. D.V.Z and K.P performed and analyzed most of *in vitro* experiments. L.L-E. performed kinome experiments. L.L-E and L.F analyzed kinome experiments. D.V.Z., W.C., K.H, M.I. and X-P.B. performed *in vivo* experiments. D.V.Z and N.Z. performed calcium imaging. J.L-A. and M.M.A. performed mitochondrial cristae analysis. C.R. performed tissue processing and histological immunofluorescence. M-A.B and J.R. performed MERC purification. G.P. performed PLA experiments. H.G-A. and J.I. performed and analyzed metabolomics experiments. D.V.Z. and L.F. wrote the manuscript. All authors approved the final version of the article.

## Acknowledgment

The authors would like to thank the lab members of Prof. Lluis Fajas group for the helpful discussions, Manfredo Quadroni and the Proteomic Analysis Facility (PAF), Arnaud Paradis and the Cellular Imaging Facility (CIF), the Genomic Technologies Facility (GTF), Jean Daraspe and Antonio Mucciolo from Electron Microscopy Facility (EMF), the FBM animal facility platform, and the Scientific Service at the Center for Integrative Genomics (SSC). This work was supported by the University of Lausanne. J.L-A. was supported by a FPI fellowship (FPI17/BES-2017-081354) from the Ministerio de Ciencia e Innovación (Spain), and a Short-Term Fellowship (8855) from the European Molecular Biology Organization (EMBO).

