## Supplementary Figures for "CDK4 inactivation balances resistance to apoptosis with heightened metabolic sensitivity in triple negative breast cancer cells"

#### Supplementary Figure 1

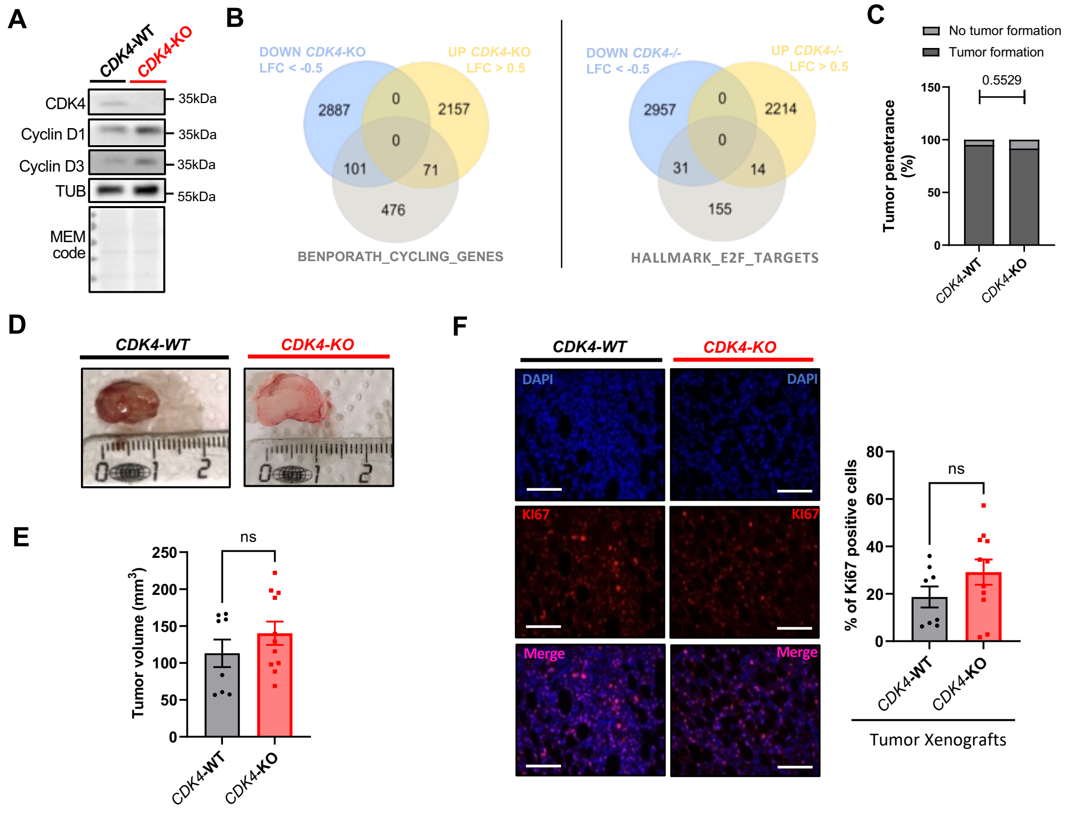

**(A).** Immunoblots and relative protein levels of CDK4, Cyclin D1, Cyclin D3 and Tubulin (TUB), of CDK4-WT and -KO TNBC cells, representative of N=6 independent biological replicates**. (B).** Venn diagram for RNA-seq data for genes differentially downregulated (log fold change < -0.5) or upregulated (log fold change > 0.5) and intersection with GSEA from cycling genes or E2F-target associated. **(C).** Tumor xenograft penetrance. N=20 WT and N=10 KO. Fisher Exact T-Test. **(D-E).** Representative tumor xenograft pictures and tumor xenograft volume at the sacrifice. N=8 WT and N=10 KO. Mann Whitney test. **(F).** Immunofluorescence of slices of CDK4-WT and -KO xenografts for KI67 (red channel) and DAPI (blue channel). Representative pictures and associated quantification of Ki67 positive cells. Scale bar: 200µm. N=8 WT and N=10 KO. Unpaired T-test. ns: non-significant.

### Supplementary Figure 2

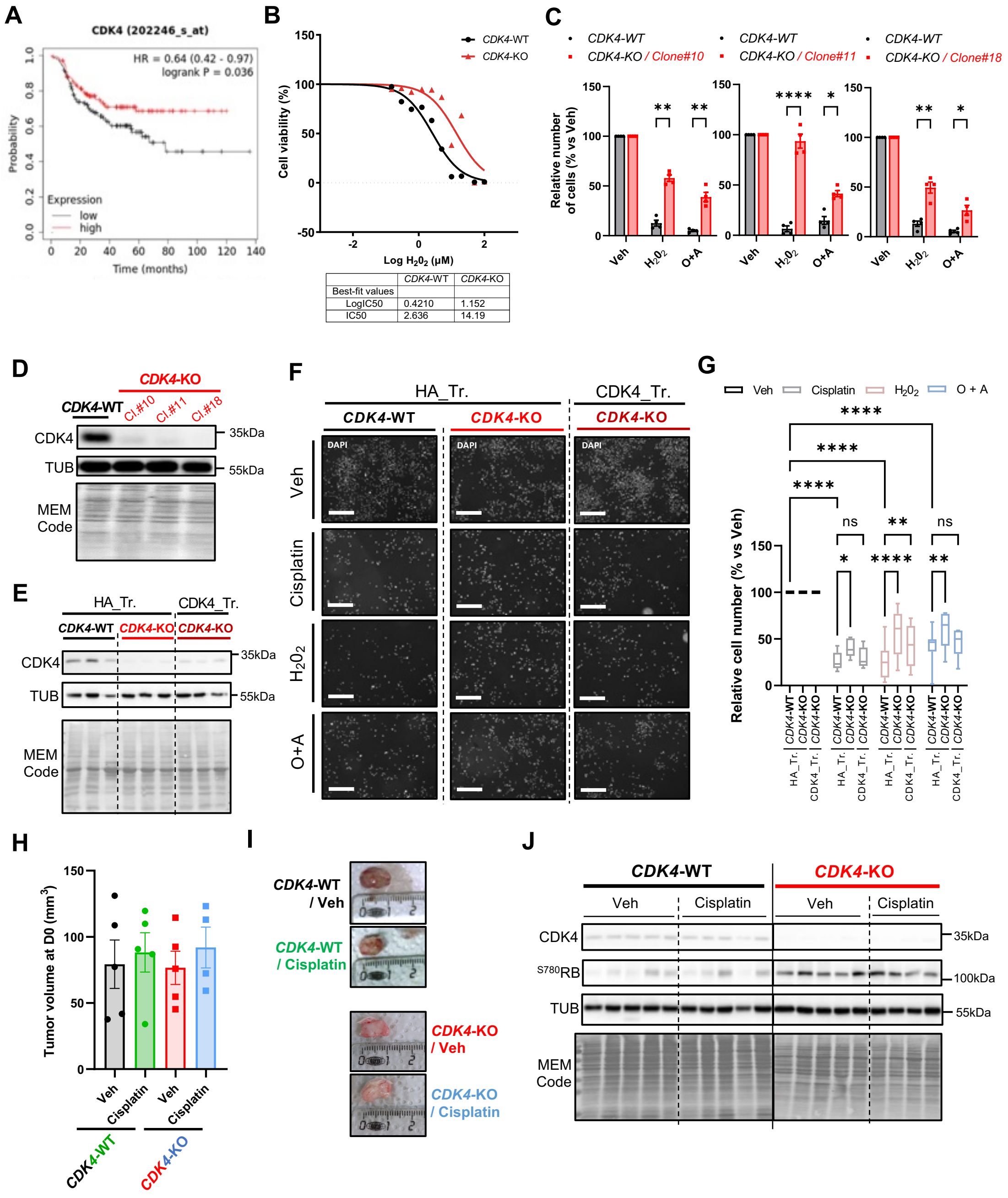

#### Supplementary Figure 2

**(A).** Kaplan-Meier Plots (prognosis curves) of patients with Hormone Receptors negative breast cancers and chemotherapy-treated, based on low (black) or high (red) CDK4 expression, N=275 patients. Source: Kmplot, from Győrffy B., Computational and Structural Biotechnology Journal, 2021;19:4101-4109, https://doi.org/10.1016/j.csbj.2021.07.014. Log-Rank T-test. **(B).** IC50 values for $H_20_2$ in cytotoxicity assay evaluating remaining viable cells 2 days after treatment. Representative of N=3 independent biological replicates. **(C).** Quantification of number of CDK4-WT and different CDK4-KO clones (#10, #11 and #18) of TNBC cells, upon treatment with Vehicle, $H_20_2$ or Oligomycin and Antimycin (O+A). N=4 independent biological replicates. 2way ANOVA and Sidák's multiple comparisons tests. **(D).** Immunoblots of CDK4, Tubulin (TUB) and MEM code number of CDK4-WT and different CDK4-KO clones (#10, #11 and #18) of TNBC cells. **(E).** Immunoblots and relative protein levels of CDK4, Tubulin and MEM code of number of CDK4-WT and different CDK4-KO clones (#10, #11 and #18) of TNBC cells. Mean +/- N=3 independent biological replicates. **(F-G).** Representative pictures of DAPI staining and quantification of number of CDK4-WT, CDK4-KO TNBC cells transfected with empty plasmid HA (HA_Tr.) and CDK4-KO TNBC cells expressing endogenous CDK4 (CDK4_Tr.), upon treatment with Vehicle, Cisplatin, $H_20_2$, or Oligomycin and Antimycin (O+A). Scale bar: 400µm. N=5 independent biological replicates. Mixed-Effect Analysis and Tukey's multiple comparisons test **(H).** Tumor volume of CDK4-WT and -KO tumor xenografts after randomization and before the start of Cisplatin treatment. 2way ANOVA and Sidák's multiple comparisons tests**.** **(I).** Representative CDK4-WT and -KO tumor xenografts pictures at the end of Cisplatin treatment. **(J).** Immunoblots of CDK4, $^{S780}$RB, Tubulin (TUB) and MEM code of CDK4-WT and -KO tumor xenografts, treated either with Vehicle or Cisplatin. n=5 WT Veh, N=5 WT Cisplatin, N=5 KO Veh and N=4 KO Cisplatin. ns: non-significant/*p-value* > 0.1,* *p-value* < 0.05, ** *p-value* < 0.01, *** *p-value* < 0.001. **** *p-value* < 0.0001.

### Supplementary Figure 3

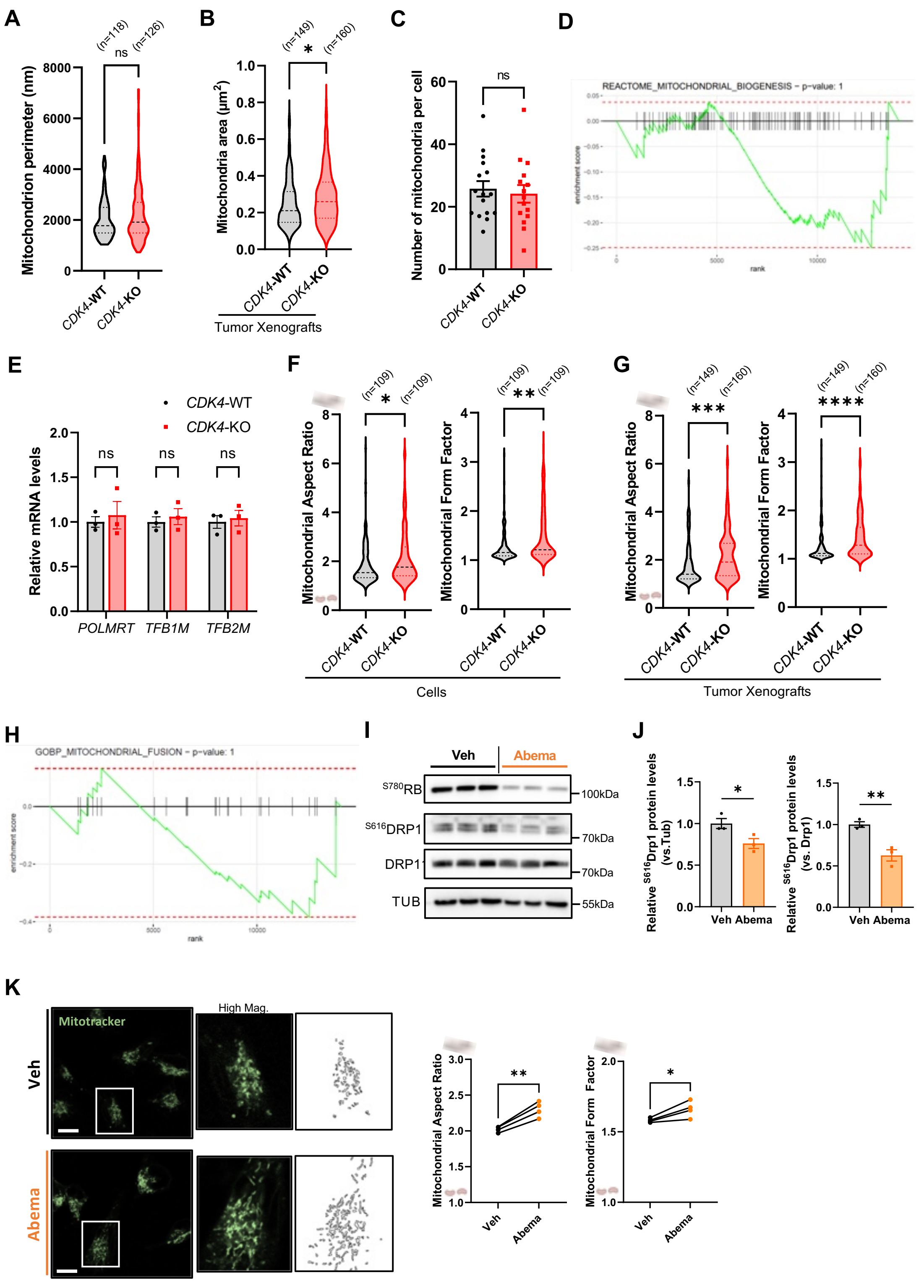

#### Supplementary Figure 3

**(A).** Mitochondria perimeter of CDK4-WT and -KO TNBC cells according to electron micrographs. n=mitochondria accounting for N=3 independent biological replicates. Mann-Whitney Test **(B).** Quantification of mitochondria area of CDK4-WT and -KO tumor xenografts. n=mitochondria number representative from 30 (CDK4-WT) and 41 (CDK4-KO) cells of N=2 tumor xenografts. Mann-Whitney Test. **(C).** Mitochondrial number per cell of CDK4-WT and -KO TNBC cells according to electron micrographs (n=16 CDK4 +/+ cells and n=15 CDK4 -/- cells). Unpaired T-test. **(D).** GSEA Analysis for REACTOME_MITOCHONDRIAL_BIOGENESIS dataset from RNA-seq data on CDK4-WT and -KO TNBC cells. **(E).** Relative mRNA levels of *POLMRT*, *TFB1M* and *TFB2M* genes of CDK4-WT and -KO TNBC cells. N=3 independent biological replicates. Multiple unpaired T-tests. **(F).** Mitochondrial aspect ratio (major axis/minor axis) and form factor (1/circularity) of CDK4-WT and -KO TNBC cells according to electron micrographs. n=mitochondria number representative from N=3 independent biological replicates. Mann-Whitney Test. **(G).** Mitochondrial aspect ratio and form factor of CDK4-WT and -KO tumors xenografts. n=mitochondria number representative from 30 (CDK4-WT) and 41 (CDK4-KO) cells of N=2 tumor xenografts. **(H).** GSEA Analysis for GOBP_MITOCHONDRIAL_FUSION dataset from RNA-seq data on CDK4-WT and -KO TNBC cells. **(I-J).** Immunoblots and relative protein levels of $^{S780}$RB, $^{S616}$DRP1, DRP1 and Tubulin (TUB) of TBNC cells pre-treated cells with Vehicle or Abemaciclib (Abema) for 8 days. N=3 independent biological replicates. Unpaired T-tests. **(K).** Representative pictures of mitochondria staining using Mitotracker of TBNC cells pre-treated cells with Vehicle or Abemaciclib (Abema) for 8 days. Associated quantification of mitochondrial aspect ratio and form factor. N=4 independent biological replicates representing of n=23 (CDK4-WT) and n=20 (CDK4-KO) cells. Paired T-tests. ns: non-significant/*p-value* > 0.1,* *p-value* < 0.05, ** *p-value* < 0.01, *** *p-value* < 0.001.

#### Supplementary Figure 4

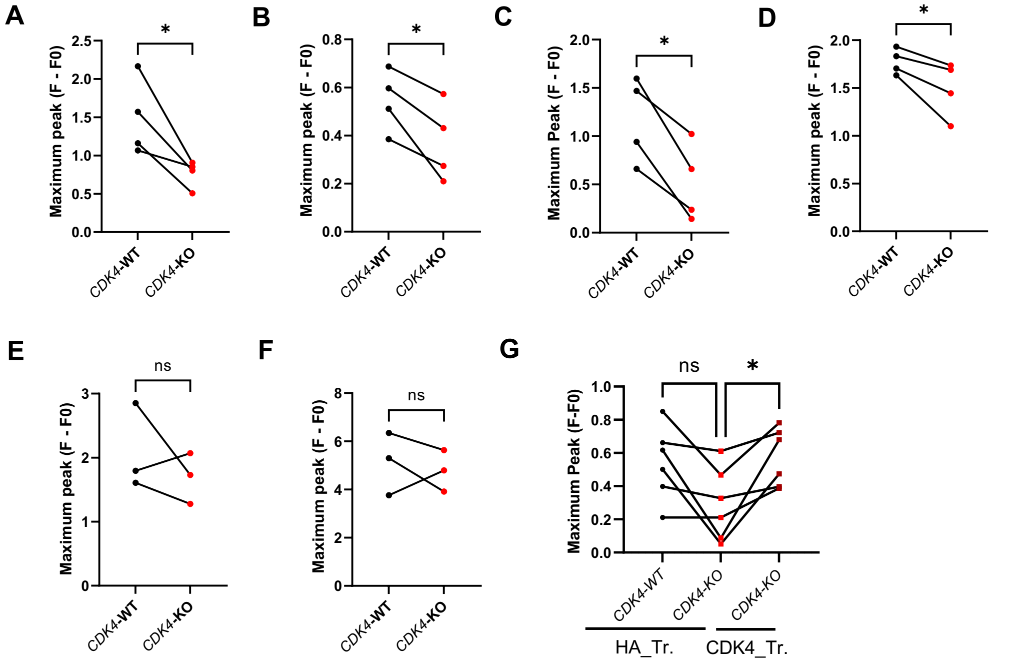

**(A-D).** Maximum amplitude/peak from basal mitochondrial calcium levels upon $H_20_2$ (2,5mM) or Oligomycin+Antimycin A (O+A) (100 µM, 10µM), Thapsigargin (TG) (2µM) or Histamine (Hist) (50 µM) injections injections. N=4 independent biological replicates representing a total of n=135 cells for 10 independent injections (WT-$H_20_2$), n=123 cells for 9 independent injections (KO-$H_20_2$), n=128 cells for 9 independent injections (WT-O+A), n=128 cells for 9 independent injctions (KO-O+A), n=148 cells for 13 independent injections (WT-TG), n=142 cells for 13 independent injections (KO-TG), n=133 cells for 10 independent injections (WT-Hist.) and n=141 cells for 12 independent injections (KO-Hist.). Paired T-tests. **(E-F).** Maximum amplitude/peak from basal cytosolic calcium levels upon Thapsigargin (TG) (2µM) or Histamine (Hist) (50 µM) injections. N=3 independent biological replicates representing a total of n=107 cells for 9 independent injections (WT-TG), n=123 cells for 9 independent injections (KO-TG), n=121 cells for 9 independent injections (WT-Hist.) and n=118 cells for 9 independent injections (KO-Hist.). Paired T-tests. **(F).** Maximum amplitude/peak from basal mitochondrial calcium levels of CDK4-WT, CDK4-KO TNBC cells transfected with empty plasmid HA (HA_Tr.) and CDK4-KO TNBC cells expressing endogenous CDK4 (CDK4_Tr.) upon Thapsigargin (TG) treatment. N=6 independent injections accounting 2 biological replicates representing a total of n=75 cells (WT/HA_Tr.), n=55 cells (KO/HA_Tr.) and n=75 cells (KO/CDK4_Tr.). RM 1way ANOVA and Tukey's multiple comparisons test. ns: non-significant/*p-value* > 0.1,* *p-value* < 0.05.

#### Supplementary Figure 5

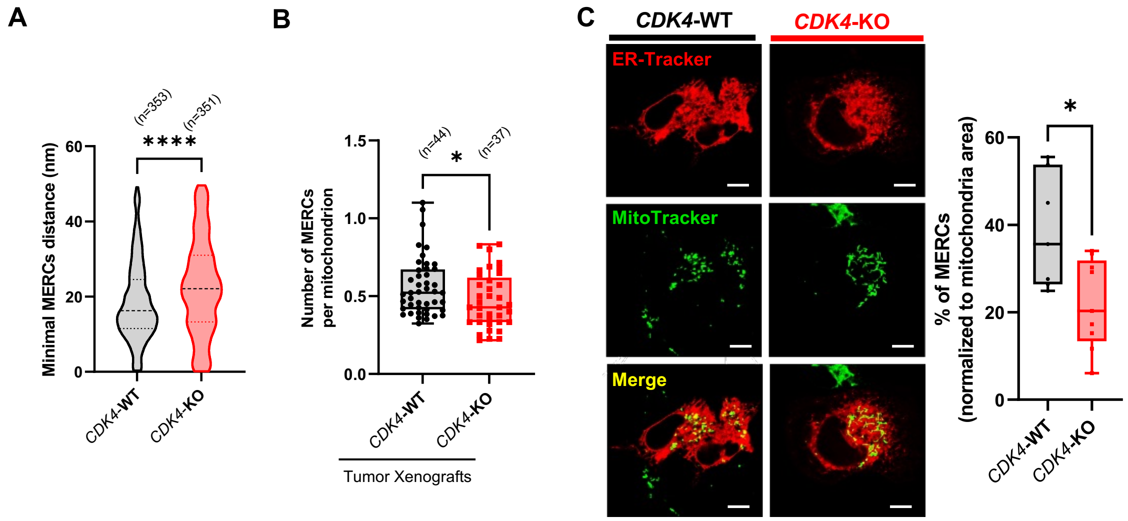

**(A).** Minimum ER-mitochondria distance in analyzed MERCs**.** N=3 independent biological replicates representing n(WT)=353 MERCs and n(KO)=351 MERCs. Mann-Whitney Test. **(B).** Quantification of number of MERCs per mitochondrion of CDK4-WT and CDK4-KO xenografts. N=2 independent xenograft samples representing a total of n(WT)=44 cells and n(KO)=37 cells. Mann-Whitney Test. **(C).** Representative micrographs of CDK4-WT and CDK4-KO TNBC cells labelled with ER tracker (red) and MitoTracker (green). Scale bar: 10 µm Quantification of colocalization MERCs, normalized to Mitochondria Area, as indicated by % of MERCs. n(WT)=7 cells and n(KO)=9 cells. * *p-value* < 0.05, **** *p-value* < 0.0001.

**Supplementary Figure 6**

**A**

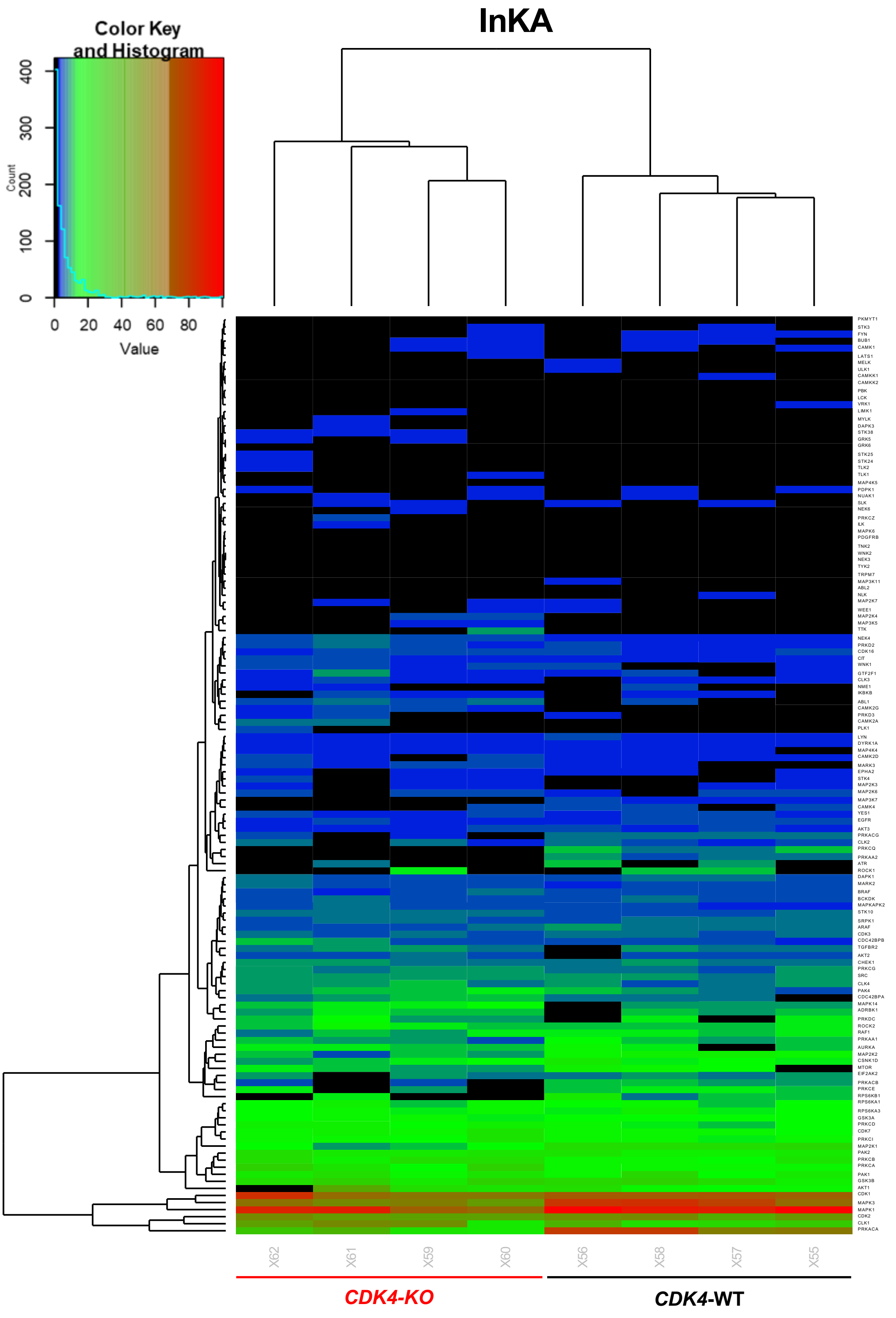

#### Supplementary Figure 6

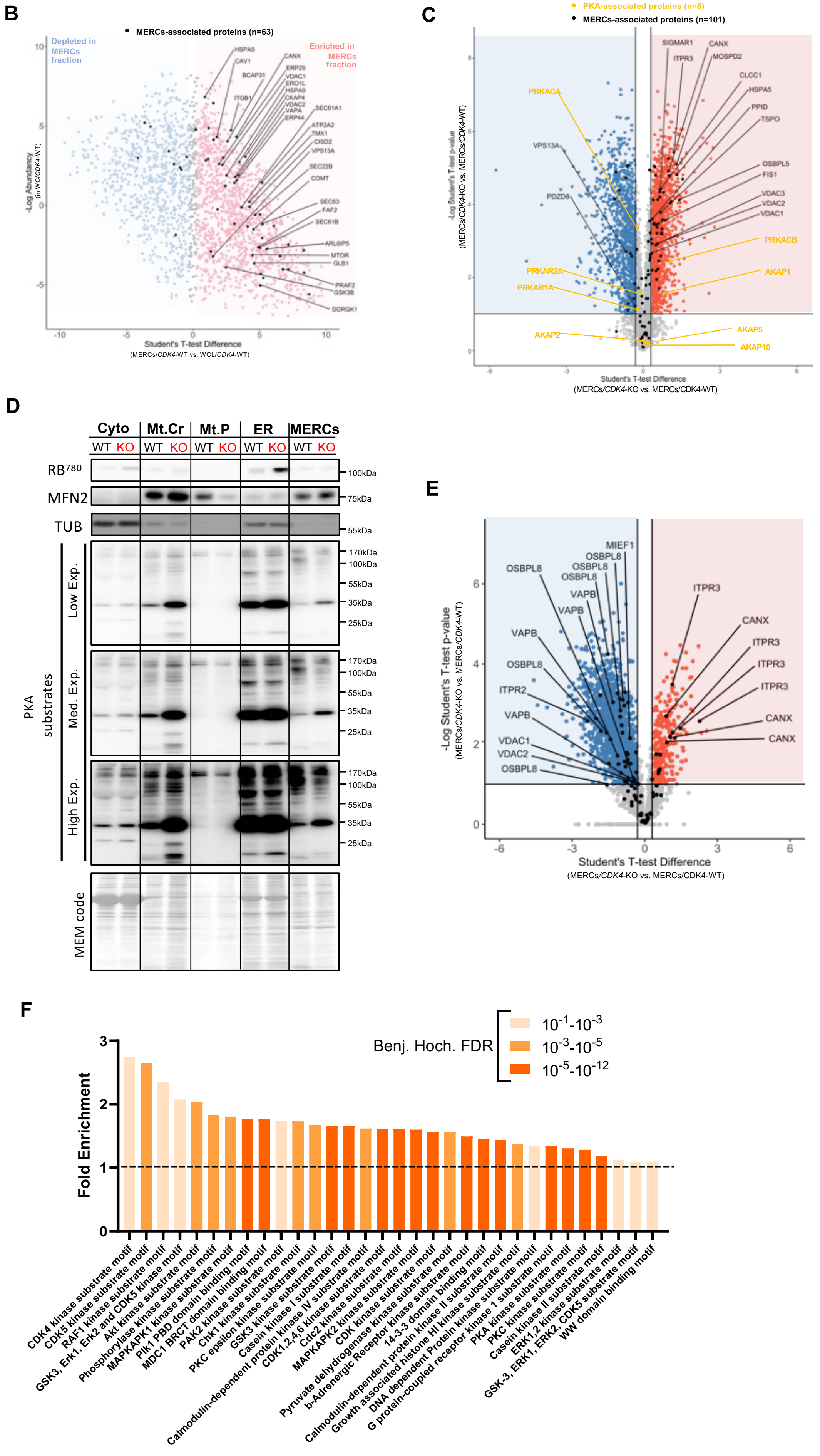

#### Supplementary Figure 6

**(A).** Heatmap of InKA analysis for kinases activity in CDK4-WT and CDK4-KO TNBC cells. N=4 independent biological replicates. **(B).** Plot representing relative abundancy of proteins detected though LC-MS/MS analyses in both whole cell lysate (WCL) and MERCs fractions of CDK4-WT TNBC cells. Student's T-test difference displays relative proteins enrichment in MERCs fractions of CDK4-WT TNBC cells. **(C).** Volcano plot of proteomics data from the MERCs fraction of CDK4-WT and CDK4-KO TNBC cells. **(D).** Immunoblots of $^{S780}$RB, MFN2, Tubulin (TUB), Phospho-PKA substrates and MEM code of CDK4-WT and CDK4-KO TNBC cells on different subcellular fractions: whole cell lysate, cytosolic, crude mitochondria, pure mitochondria, ER and Mitochondria-ER contacts. Representative of N=3 biological replicates. **(E).** Volcano plot of phospho-proteomics data from the MERCs fraction of CDK4-WT and CDK4-KO TNBC cells. **(F).** Enrichment analysis on phospho-peptides found in MERCS fraction of CDK4-WT and -KO TNBC cells. Benj. Hoch. FDR value ranges are displayed.

### Supplementary Table 1

| Protein Name | Main Associated-Function |
|---|---|
| BCL2 | Apoptosis |
| MCL1 | |
| BECN1 | |
| BAX | |
| BAP1 | |
| BCL2L1 | |
| BAK1 | |
| PML | |
| BCL2L13 | |
| FUNDC2 | Autophagy |
| ATG14 | |
| STX17 | |
| GLB1 | |
| MTOR | |
| RICTOR | |
| WFS1 | ER Homeostasis |
| ERP29 | |
| HSP90B1 | |
| TMX1 | |
| DDRGK1 | |
| SRPRB | |
| SEC63 | |
| SEC22B | |
| SEC61B | |
| SEC61A1 | |
| EIF2AK3 | |
| RPS7 | |
| RTN4 | |
| ATF6 | |
| ERP44 | |
| SACM1L | |
| UBXN4 | |
| HSPA5 | |
| ITPR2 | Ions Transport |
| ATP2A2 | |
| RYR3 | |
| MCU | |
| VDAC3 | |
| VDAC1 | |
| VDAC2 | |
| CANX | |
| CLCC1 | |
| ITPR3 | |
| TSPO | Lipids |
| HSD17B12 | |
| DHCR7 | |
| HK2 | Mitochondrial Bioenergetics |
| SLC25A5 | |
| GSK3B | |
| CYC1 | |
| CYCS | |
| CYB5R3 | |
| NDUFA8 | |
| KRAS | Proliferation |
| AKT3 | |
| ANXA7 | |
| PPP2CA | |
| PTEN | |
| AKT2 | |
| TP53 | |
| AKT1 | |
| CAV1 | |

| Protein Name | Main Associated-Function |
|---|---|
| ARMCX3 | Others |
| PGRMC1 | |
| TCHP | |
| CKAP4 | |
| NAPG | |
| OCIAD1 | |
| VPS13A | |
| FUS | |
| PGRMC2 | |
| VAPA | |
| CCDC47 | |
| HNRNPR | |
| TGM2 | |
| EXD2 | |
| SCCPDH | |
| TARDBP | |
| EEF1D | |
| TDRKH | |
| RAB1A | |
| COMT | |
| TOMM70A | |
| PARK7 | |
| SAR1B | |
| LRRC59 | |
| PTRH2 | |
| PRAF2 | |
| ITGB1 | |
| SAR1A | |
| TBL2 | |
| ARL6IP5 | |
| CPD | |
| MAVS | |
| PSEN1 | |
| MTX2 | |
| STT3B | |
| HMOX2 | |
| INF2 | |
| AHCYL1 | |
| RHOT1 | |
| FAF2 | |
| APP | |
| VPS13C | |
| MIEF1 | Mitochondrial Dynamics |
| MFN2 | |
| MFF | |
| MIEF2 | |
| MFN1 | |
| ERO1L | Redox Signaling |
| CISD2 | Tetheriig |
| BCAP31 | |
| OSBPL8 | |
| FKBP8 | |
| HSPA9 | |
| ATAD3A | |
| PDZD8 | |
| PPID | |
| FIS1 | |
| VAPB | |
| SIGMAR1 | |
| RMDN3 | |
| OSBPL5 | |
| MOSPD2 | |

#### Supplementary Table 2

| Gene | Sequence |
|---|---|
| ***Human*** | |
| *RS9*-Forward | CACACTCTCCCCAACGTTCT |
| *RS9*-Reverse | ACCACCTGCTTGCCGGACCCTGA |
| *TBP*-Forward | TGCACAGGAGCCAAGAGTGAA |
| *TBP*-Reverse | CACATCACAGCTCCCCACCA |
| *POLMRT*-Forward | ACGGAGACATCAGCCGAAAG |
| *POLMRT*-Reverse | GGCCCTTCCTGTAGCAGTG |
| *TFB1M*-Forward | CGGAAAACTCAGCACTTGCC |
| *TFB1M*-Reverse | ACAATCTTATCTGTCAGCCTCAAGT |
| *TFB2M*-Forward | ATGTCTTCTCGAGGGCTCTTT |
| *TFB2M*-Reverse | TTAAAGGGATGTCTGCTGTCCAA |

#### Supplementary Table 3

| Protein | Reference | Dilution |
|---|---|---|
| CDK4 | #1790 (D9G3E) (Cell Signaling Technology) | 1/1000 |
| CDK6 | #3136 (Cell Signaling Technology) | 1/2000 |
| Cyclin D1 | PA5-16607 (ThermoFisher) | 1/250 |
| Cyclin D3 | ab28283 [DCS2.2] (Abcam) | 1/1000 |
| $^{S780}$RB | #8180 (Cell Signaling Technology) | 1/1000 |
| RB | sc-50 (SantaCruz) | 1/1000 |
| Cleaved Casp-3 | #9664 (Cell Signaling Technology) | 1/1000 |
| $^{S616}$DRP1 | #4494 (D9A1) (Cell Signaling Technology) | 1/1000 |
| DRP1 | #5391 (D8H5) (Cell Signaling Technology) | 1/1000 |
| SERCA1 | Ab129104 [EPR7322] (Abcam) | 1/1000 |
| $^{S1756}$ITPR1 | #3760 (Cell Signaling) | 1/1000 |
| ITPR1 | 07-1213 (MERCK) Lot#3546915 | 1/1000 |
| ITPR2 | sc-398434 (Santa Cruz) | 1/1000 |
| ITPR3 | 610313 (BD Tr.) Lot#2/IP3R-3 | 1/1000 |
| VDAC1 | ab14734 (Abcam) Lot[20B12AF2] | 1/1000 |
| MCU | HPA016480-100UL (Sigma) | 1/1000 |
| PKA Phospho-substrates | #9624 (100G7E) (Cell Signaling Technology) | 1/1000 |
| CALR | #12238 (D3E6) XP® (Cell Signaling Technology) | 1/1000 |
| PDH | A b110334 (Abcam) | 1/1000 |
| GRP75 | #2816 (Cell Signaling Technology) | 1/1000 |
| MFN2 | #11925 (D1E9) (Cell Signaling Technology) | 1/1000 |
| PRKAR1A | #5675 (D54D9) (Cell Signaling Technology) | 1/1000 |
| TUB | T6199-200UL (Sigma) | 1/5000 |
